# Equine dermatitis outbreak associated with parapoxvirus

**DOI:** 10.1101/2023.09.01.555671

**Authors:** Jenni Virtanen, Katja Hautala, Mira Utriainen, Lara Dutra, Katarina Eskola, Niina Airas, Ruut Uusitalo, Ella Ahvenainen, Teemu Smura, Tarja Sironen, Olli Vapalahti, Ravi Kant, Anna-Maija K. Virtala, Paula M. Kinnunen

## Abstract

Parapoxviruses (PPV) cause skin and mucous membrane lesions in several animal species, and of the five recognized PPVs, at least three are zoonotic. Equine PPV (EqPPV) is the sixth one initially described in humans in the United States and later in a severely sick horse in Finland in 2013–2015. In 2021–2022, a large-scale pustulo-vesicular pastern dermatitis outbreak occurred in horses all over Finland. This study aimed at analysing the outbreak, identifying and describing the causative agent, describing clinical signs, and searching for risk factors. EqPPV was identified as a probable causative agent and co-infections with several potentially pathogenic and zoonotic bacteria were observed. Histopathologically, suppurative and ulcerative dermatitis was diagnosed. Due to the lack of specific tests for this virus, we developed a novel diagnostic EqPPV-PCR with sensitivity of 10 copies/reaction. Based on a large proportion of the genome sequenced directly from clinical samples, very little variation was detected between the sequences of the case from 2013 and the cases from 2021–2022. Based on an epidemiological survey, the main risk factor for pastern dermatitis was having racehorses. Approximately one third of the horses at each affected stable got clinical dermatitis, manifesting as severe skin lesions. Skin lesions were also occasionally reported in humans, indicating potential zoonotic transmission. Case stables commonly reported attendance in race events before acquiring the disease. Survey also identified differences in practises between case and control stables. Taken together, these results enable a better preparedness, diagnostics, and guidelines for future outbreaks.

## Introduction

Poxviruses cause skin and general infections in humans and vertebrate animals (1). Parapoxviruses (PPVs) are known to cause skin and mucous membrane infections in ruminants and occasionally other animal species (2). Until recently, five virus species have been known: orf virus (ORFV), bovine papular stomatitis virus (BPSV), pseudocowpox virus (PCPV), red deer parapoxvirus (RDPV) and grey sealpox virus (GSEPV) (3). ORFV, BPSV, and PCPV are zoonotic and the rest are potentially zoonotic (3, 4, 5). In Finland, ORFV infections are detected annually in sheep, and PCPV and BPSV in bovines. Also, reindeer papular stomatitis caused by ORFV and PCPV, and their zoonotic forms are reported (6, 7).

Parapoxviruses proliferate in the corneum cells of the epidermis, resulting in pustular lesions at the site of infection (2, 8, 9). The primary lesions can be severe, but without secondary bacterial infection heal in 4–6 weeks. The secondary lesions can ulcerate and necrotize, delaying healing. In humans, infection typically manifests as vesicles on the hands after contact with infected ruminants.

In 2015, a symptomatic poxvirus infection was reported in two humans in the USA, both of whom had engaged with horses and donkeys (10). In 2013, our research team showed a poxvirus in a severely ill horse bred in Finland. The partial RNA polymerase gene was 99-100% identical on nucletiode level to the one found in human patients in the United States (11), and most similar to parapoxviruses. Almost complete genome sequencing later confirmed the virus to be a novel PPV species, Equine parapoxvirus (EqPPV) (12). However, the virus was distant from other PPVs and could not be detected by many existing PPV-PCR assays (11). No new cases were reported until December 2021, when a pastern dermatitis epidemic started in racehorses in Finland. Owners, trainers, and veterinarians estimated that hundreds of horses got sick, main symptoms being painful skin lesions in the pastern area. Sick horses could not race, and many had to stop training. A preliminary analysis of partial envelope phospholipase gene (ORF011, named according to (13) throughout the manuscript) (14) confirmed the presence of a PPV 97% identical to the 2013 case in 36% (9/25) of the sampled horses (12). However, most sampled horses remained without final diagnosis, presumably due to unsuitability of the pan-PPV-PCR for this particular virus and host (12).

In this study, we used histopathology, microbiological cultures, molecular methods, and epidemiological survey to establish a validated diagnostic protocol, and to elucidate the epidemiology as well as the clinical and microbiological picture of the dermatitis outbreak, and the poxvirus found in horses.

## Materials and methods

### Sampling

Samples were taken from fully developed vesicles or other skin lesions of 26 acutely ill horses from 11 stables (Table 1) in as early stage of the disease as possible. The samples were taken by veterinarians as a part of routine diagnostics and hence, no ethical approvals were needed. Swab samples were taken into empty sterile tubes, or a tube filled with 1 ml of sterile saline. Before analyses, dry swabs were incubated in 1 ml of Dulbecco’s PBS + 0.2% bovine serum albumin overnight at 4°C. Biopsy samples were collected from six horses preferably from the edges of skin lesions using a punch or scalpel. However, due to the difficulty of sampling, mainly vesicles or centres of the ulcers were biopsied. If possible, the biopsy samples were immediately fixed in 10% neutral buffered formalin. We also took four surface swab samples (15) from a racecourse environment: Two from the concrete floor of a shared washing box, and two from the metallic locks attached to a horse’s halter in the harnessing paddock area. Most samples were stored fresh in +4°C before and during shipment (mean and median 2 days, range 0–5 days) after which they were stored in −80°C. The frozen samples were melted and fixed in formalin later, before histopathological analysis.

**Table 1.**
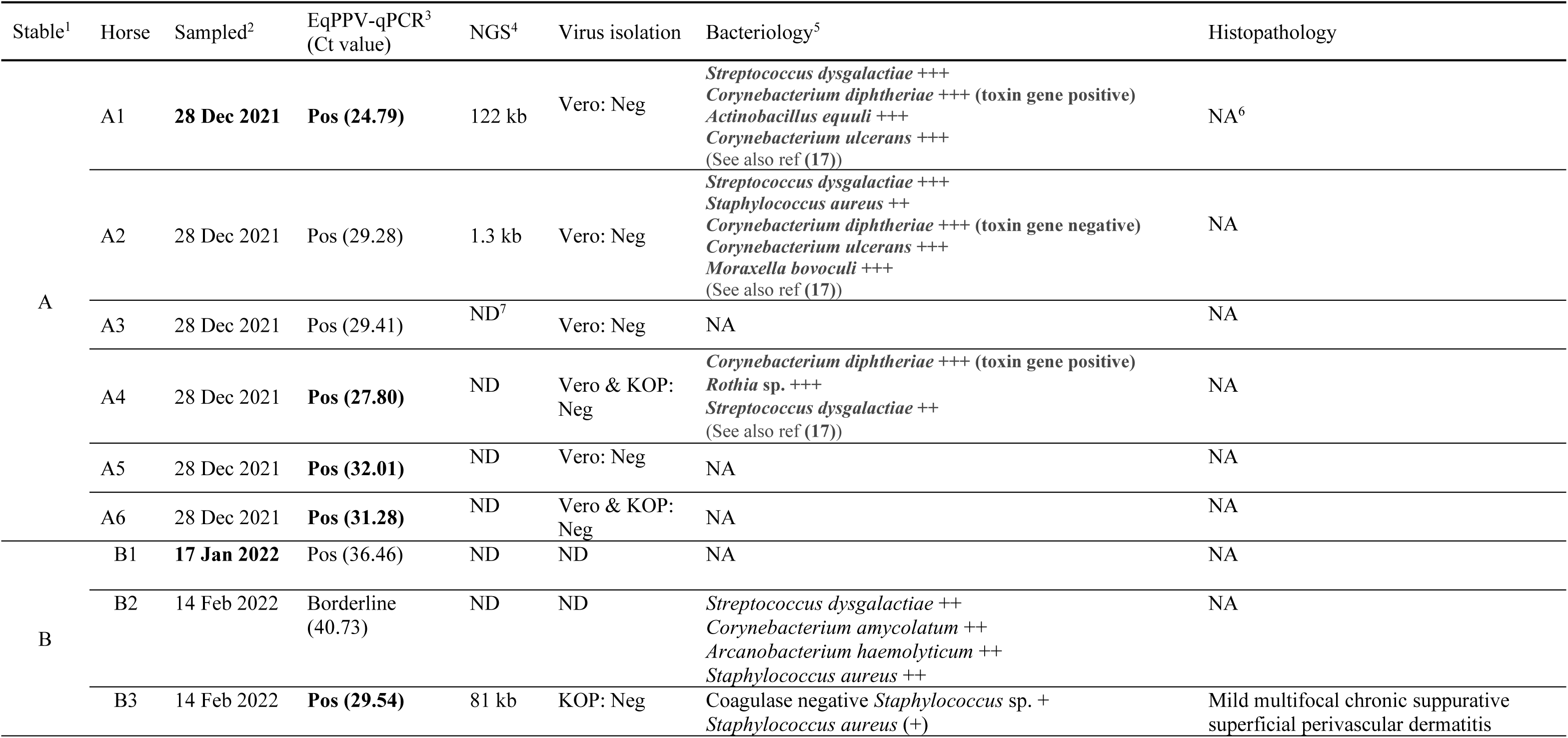
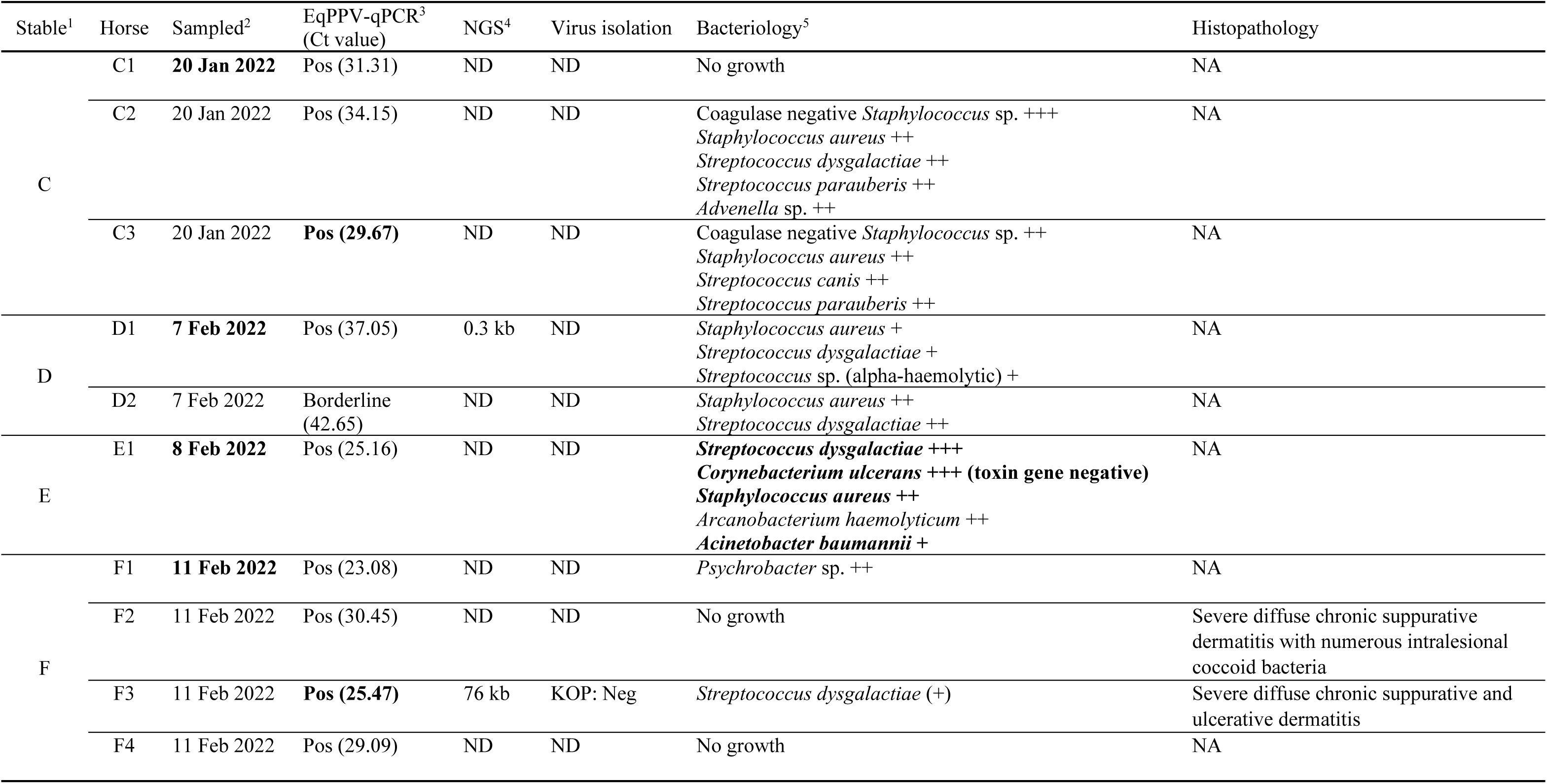
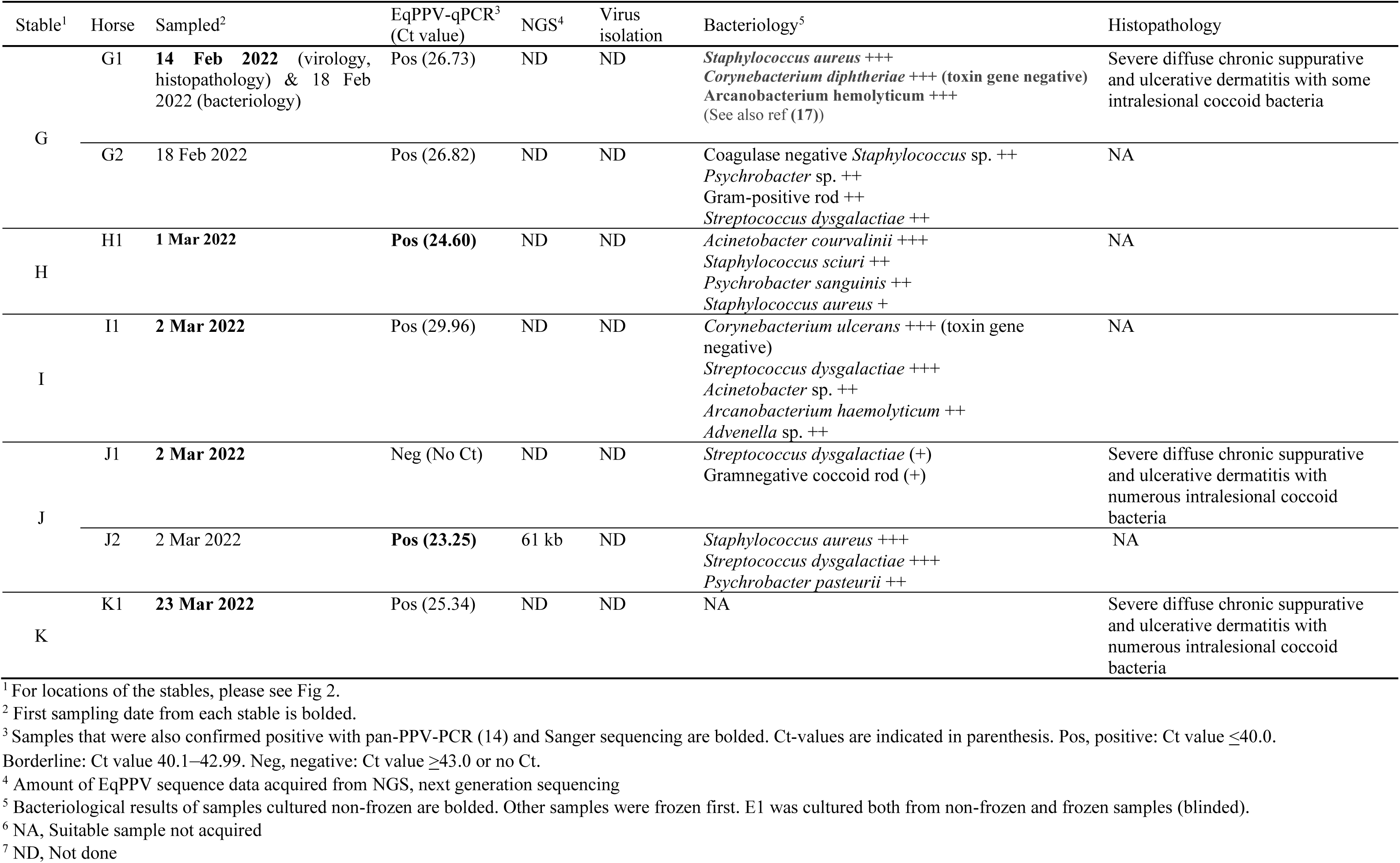
Laboratory findings of the horses with clinically diagnosed pastern dermatitis.

### Histopathology

The formalin-fixed biopsy samples were routinely processed and embedded in paraffin wax. Sections were cut at 4 µm and stained with haematoxylin and eosin for light microscopic examination. Selected samples were also stained by Gram stain and by periodic acid-Schiff. A board-certified veterinary pathologist performed histological analyses.

### Bacteriological analyses

Bacteriological culturing was performed in the Clinical Microbiology Laboratory of the Faculty of Veterinary Medicine (University of Helsinki). Five samples (A1, A2, A4, G1, E1) were cultured from a separate direct smear obtained with a cotton swab in sterile bacterial culture transport medium from the lesion sites. The frozen samples were cultured by suspending the defrosted sample in 1 ml of sterile NaCl and culturing two drops of the suspension. E1 was cultured blinded both from non-frozen and frozen samples for comparison. The culturing for all the samples was done onto tryptic soy agar with 5% sheep’s blood (Thermo Fisher Scientific, USA). The plates were incubated at +35±2°C in 5% CO_2_ and read at 24-hour intervals for four days. The phenotypically different bacterial colonies were identified using conventional bacteriological methods as well as matrix-assisted laser desorption/ionization time-of-flight mass spectrometry (Bruker MALDI Biotyper Microflex LT, Bruker Daltonik GmbH, Bremen, Germany). The amount of growth was assessed semi-quantitatively from none (-) to scarce (+) to notable (++/+++). Diptheria-toxin genes were detected with PCR from the pure cultures of selected *Corynebacterium ulcerans* isolates as described previously (16) in the Finnish Food Authority laboratory for Veterinary Bacteriology and Pathology. The toxin gene PCR for *C. diphtheriae* isolates was done as described by Grönthal et al. (17, submitted manuscript).

### Molecular methods

We used DNA extracted from swab samples previously (12) and most of the samples had already been tested with pan-PPV-PCR described by Inoshima et al (14). Six samples from five stables were subjected to next generation sequencing (NGS) with previously described protocol (12) and assembled with LazyPipe2 (18, 19). Genes were annotated with Open Reading Frame Finder (NCBI) with minimum ORF length of 150. All the annotations were checked with Blast (NCBI) and only complete protein sequences previously annotated to other PPVs were considered. Genome structure was built using Mega11, ClustalW, and UGENE (20, 21, 22).

A diagnostic qPCR for EqPPV was set up by designing primers and probe based on myristylated protein (ORF093) sequences acquired from NGS (bp 752–872). PCR reaction contained 4 µl of 5x HOT FIREPol Probe Universal qPCR Mix, 600 nm of primers EqPPV F1 (ACGAGTGCACAGACCAGTC) and EqPPV R1 (GAGTTCTCCATCACCAGGCT), 100 nm of probe EqPPV P1 (FAM-AAGTGGCTGTTCTTCAGCCAGGA-MGB), 2 µl of DNA, and water up to 20 µl. Program had a 10 min polymerase activation at 95°C followed by 45 cycles of denaturation (95°C, 15 s) and annealing/extension (58°C, 60 s). A synthetic plasmid control based on bp 740–891 of the F14.1158H (OQ110629.1, (12)) was ordered from GeneArt (ThermoFisher Scientific). Analytic sensitivity of the PCR was determined with a plasmid dilution series of 0, 1, 10, 100, and 1000 copies/reaction as the lowest copy number that was positive with all the five replicates. PCRs were performed with Stratagene Mx3005P (Agilent Technologies) with a 0.04 threshold. For clinical samples, Ct values <40 were considered positive, values 40–43 borderline, and values >43 negative. Specificity was tested with DNA from skin tissue of three healthy horses and skin swabs collected before the outbreak from two horses. All the clinical samples were tested with the established EqPPV-PCR. In addition, two clinical samples containing ORFV and PCPV DNA, were tested.

To compare a larger number of virus variants from different stables, partial sequence of ORF093 (bp 349-872) was amplified in a reaction containing 12.5 µl of Fermentas SYBR MasterMix (Thermo Scientific), 0.75 µl of primers EqPPV R1 (10 µM) and EqPPV F2 (TTCAAGATAGAGCCCGCGG), 5 µl of template, and water up to 25 µl. Program was run with Stratagene Mx3005P as follows: 95°C 10 min, 40 cycles of 95°C 30 s, 55°C 1 min and 72°C 1 min, followed by melting curve analysis at 95°C for 1 min, 55°C 30 s, and 95°C 30 s. We run products on 1.5% agarose gel, extracted bands of the correct size with Gel extraction kit (GeneJet) and sequenced with Sanger sequencing. Sequence analysis and phylogenetic tree was done with Mega11, IQ-TREE 2.2.0.7 and iTOL as previously described (12, 21, 23, 24, 25).

### Virus culture

Of eight selected PCR-positive samples, virus isolation was attempted in African Green Monkey Kidney cells (Vero E6) or primary bovine oesophagus cells (KOP) or both as described previously (12, 26). Culture supernatants showing possible cytopathic effect were tested with the qPCR described above.

### Epidemiological survey

Epidemiological survey was used to gain understanding on the epidemiology, clinical signs, treatment, distribution, risk factors, and zoonotic potential. They web survey was created in Finnish (Elomake, version 4, Eduix Ltd, https://eduix.com) and amended based on the feedback of three external testers. The survey for all respondents covered information about the stables and their animals such as number of horses, housing system, bedding, animal species, hand hygiene, the use of horses (e.g., breeding, riding, trotting), potential clinical signs in horses outside the pastern area, and skin signs in other animals or humans. Those who answered “yes” to the case definition “In the stable, one or more horses have had clinical signs of contagious pastern dermatitis (vesicles, wounds, or swelling, potentially excretion) between 1 December 2021 and 31 March 2022”, were classified as “cases” and got further series of questions. These included details of the signs and their duration and treatment, quality of the surface at outdoor enclosures, events, actions and visits during the month before the onset of outbreak at the stable, and an opportunity to express thoughts about the transmission route. Those answering “no” to the case definition question were classified as controls and got some similar but not identical questions as case stables, namely events, actions, and visits between 1 December 2021 and 31 March 2022. In all the questions, “horses” also included ponies.

The survey was available from 23 June to 31 August 2022. Link to the survey was primarily shared in social media, at first in equine-related Facebook pages “TerveTalli” and “TalkRavi”, from where it was further shared using snowball sampling (27, 28) by readers in Facebook, Twitter, and LinkedIn. Also, two newspapers published the link in their web pages. All the respondents gave their informed consent before answering the survey. We did not ask names and specifically forbid reporting any sensitive patient information. Still, we processed the collected data in accordance with the General Data Protection Regulation 2016/679 of the European Parliament and the Council. The instructions emphasized that only one response from each stable was needed.

### Statistical analysis of the survey data

Statistical analyses were carried out using IBM SPSS Statistics for Windows, version 29.0.1 (Armonk, NY: IBM Corp, USA). Descriptive data collected from case stables were described using frequency tables. Case and control stables were compared using cross-tabulation with Chi-Square tests and for non-categorical data Independent-Samples Mann-Whitney U test. Additionally, we compared stables that had at least one racehorse with those stables that had none using same explanatory variables and tests as for case-control comparisons. This was done to check for possible confounding effect of this division of the stables. We considered statistical significance at 5% alpha-level. The 95% confidence intervals for proportions were calculated using Wilson’s method (29) with an online tool (http://epitools.ausvet.com.au/content.php?page=CIProportion). The obtained p-values were adjusted for multiple comparisons by Benjamini-Hochberg correction (30) using the False Discovery Rate calculator (https://www.sdmproject.com/utilities/?show=FDR).

### Mapping

The map was created using ESRI ArcGIS (version 10.3.1) (ESRI, Redlands, CA, USA). Postal code areas (31) were downloaded through the interface service. They were merged to correspond to the general regions of postal code areas which comprise the first and second digits of five-digit numeric postal codes (32).

## Results

### The disease led to potentially long-lasting clinical signs and possible zoonotic transmission

Our group was first contacted by veterinarians and trainers about a new equine pastern dermatitis in December 2021, and numerous contacts continued until March 2022. Clinical signs typically started with pustular-vesicular lesions (Fig. 1A-B and 1D) and leg swelling. Later, the skin lesions presented with serous (Fig. 1C) or purulent excretion. Due to the pain, the horses became lame even when walking, and many contracted secondary bacterial infections (Fig. 1E). They usually recovered from the skin signs within a few weeks, but long-lasting hoof lesions were also reported (Fig. 1F). To identify and characterize the cause, practitioners sent skin swabs for molecular and bacteriological diagnostics (Table 1 and Fig. 2) and additional skin samples for histopathological diagnostics. During the epidemic, a few people taking care of the sick horses contacted us and reported skin lesions in themselves too, but unfortunately, we did not obtain samples from them.

**FIG 1.**
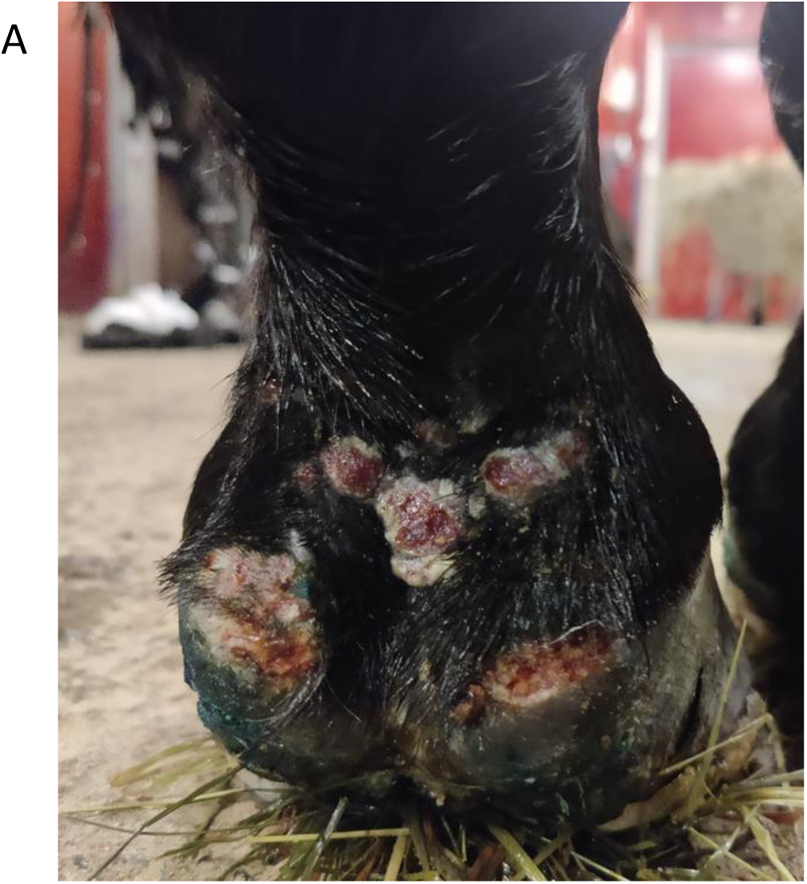
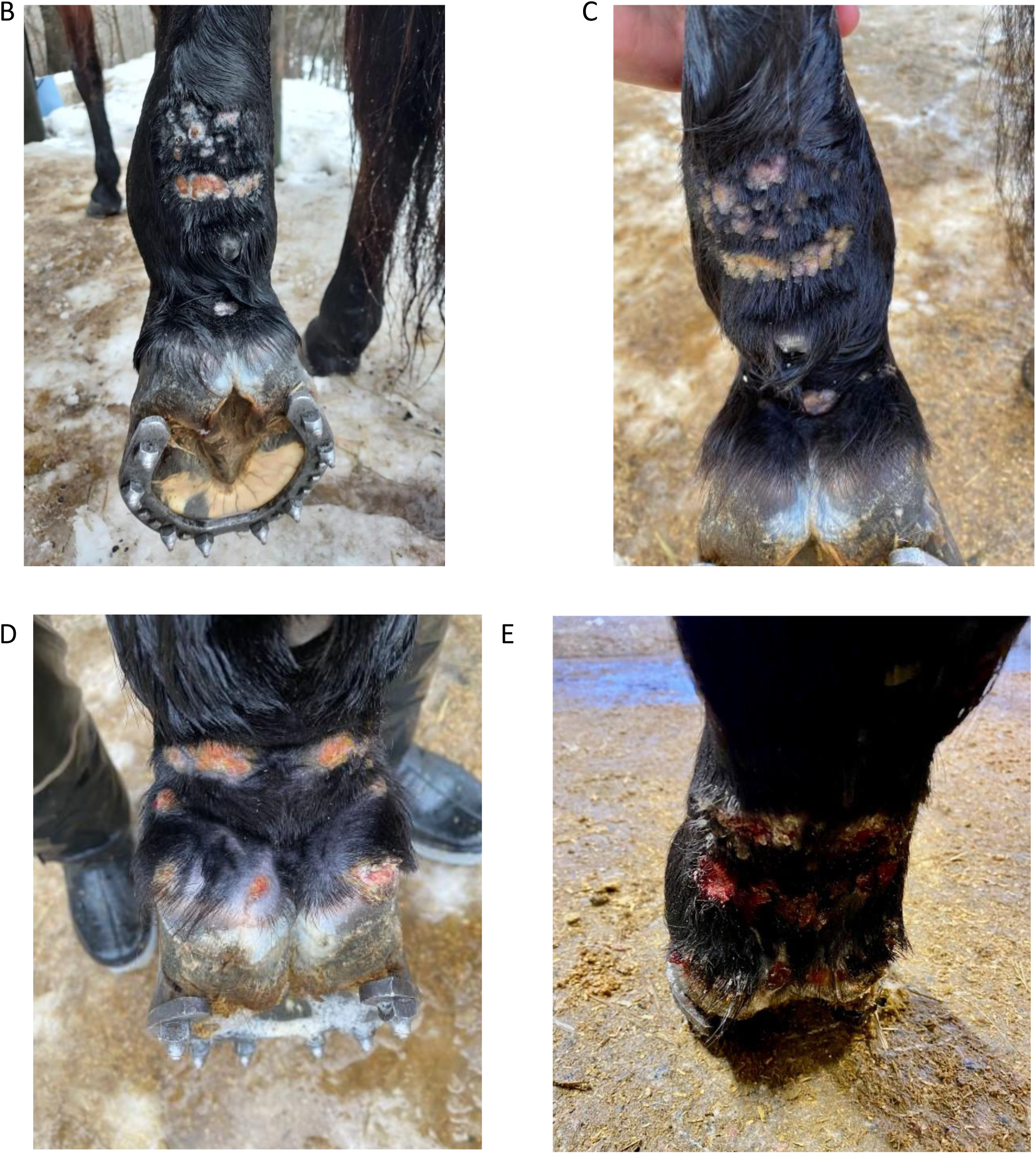
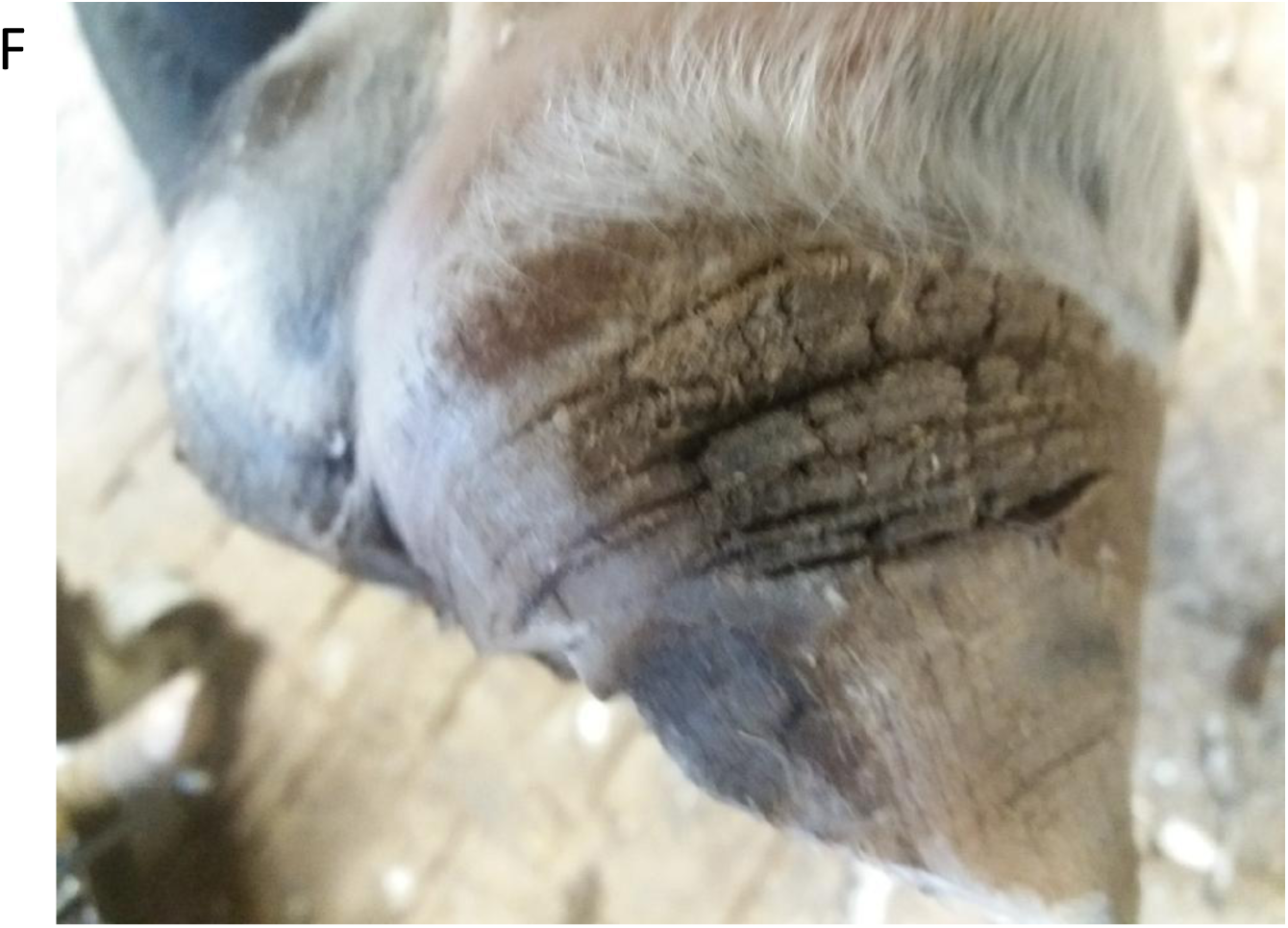
Clinical presentation of pastern dermatitis. A: Typical lesions in the pastern area (unknown horse). B and C: Pustular and vesicular primary lesions in the pastern area on horse G2 one (B) and four (C) days after the onset of clinical signs. D and E: Primary and secondary lesions on horse G1. The horse expressed first clinical signs four days after (D) training on the racetrack. On the second day after the onset of clinical signs, skin punch biopsy (4 mm) and swab samples were taken from which equine poxvirus DNA and suppurative and ulcerative dermatitis with coccoid bacteria were verified (Table 1). During the following days the clinical signs got more severe despite the local treatments, and the leg was very oedematous and painful. Six days after the onset of clinical signs, intra-muscular penicillin was initiated. After five days of penicillin treatment, secondary bacterial infection was still obvious (E) and bacterial culture showed penicillin G resistant *Staphylococcus aureus*, penicillin G sensitive *Corynebacterium diphtheriae* and *Arcanobacterium hemolyticum*. F: Chronic hoof lesions two months after the onset of severely oedematous pastern dermatitis (unknown horse).

**FIG 2.**
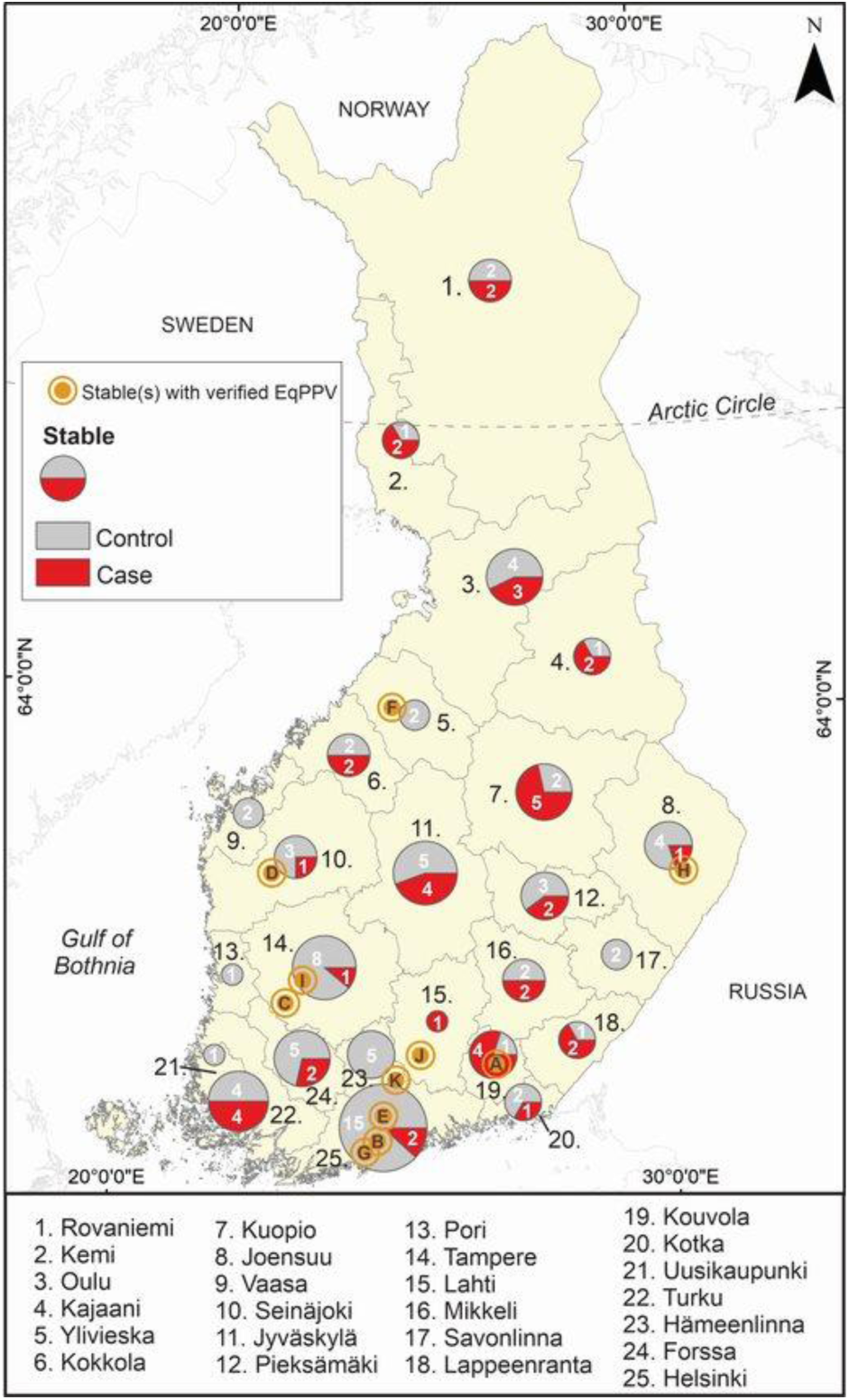
Map showing stables with PCR-confirmed EqPPV (orange symbol; Table 1) and numbers of control and case stables (pie chart) included in the study. The names with black colour indicate the general areas of post code areas.

### Severe suppurative and ulcerative dermatitis was detected

Skin biopsies consisting of epidermis and superficial dermis obtained from sick horses were subjected to histopathologic examination. The epidermis was mainly lost (ulcer) and covered by thick serocellular crust composed of cellular debris, keratin, and fibrin in most of the samples (Table 1). Numerous neutrophilic granulocytes and coccoid bacteria were embedded in crusts and seen in superficial dermis with fewer lymphocytes and plasma cells. In one sample obtained outside the ulcer, epidermis was intact and only mild population of perivascular neutrophilic granulocytes, lymphocytes and plasma cells were seen in the superficial dermis (Table 1).

### Potentially pathogenic bacteria were detected in lesions

Altogether 22 samples were obtained for bacterial culture from 21 horses (Table 1). The five samples that were not frozen before culturing harboured notable growth of mixed bacterial species, and samples cultured after freezing showed bacterial growth in 14 of the 17 samples (82.4%). Of these, 85.7% (12/14) consisted of two or more bacterial species, and 14.3% (2/14) had pure growth of a single species. In total, 17 lesions (81.0%) harboured *Streptococcus* spp. and 16 (76.2%) *Staphylococcus* spp. One frozen sample had significant growth of *Arcanobacterium haemolyticum*, which was not detected before freezing; otherwise the bacterial growth was exactly the same from fresh and frozen samples. *Corynebacterium diphtheriae* was present in lesions of four horses, which was also seen in a parallel study (17, submitted manuscript) (19.0%), and *C. ulcerans* in four (19.0%). Two *C. diphtheriae* isolates were PCR-positive for toxA gene, the other two negative. The toxin PCR for two of the *C. ulcerans* isolates was negative.

### EqPPV was detected in majority of the samples with a new diagnostic PCR

Based on our earlier study with pan-PPV-PCR (12, 14), 9/25 of the skin swabs received by then were positive for EqPPV, but the rest were without a reliable diagnosis due to poor specificity and sensitivity (12). Therefore, a new diagnostic PCR was developed based on ORF093 of the equine case from 2013 (F14.1158H) (12) and four cases from 2022. In total, 23/26 (88.4 %) samples from horses with pastern dermatitis were positive with EqPPV-PCR, and from all the stables, at least one horse was positive (Table 1, Fig. 2). Analytic sensitivity of EqPPV-PCR was 10 copies/reaction (Table 2). All the five samples from healthy horses as well as ORFV and PCPV samples were negative. Samples from six PCR-positive horses were subjected to virus isolation trials. Possible cytopathic effect was detected in one sample cultured in Vero E6 cells but was not caused by EqPPV based on the high Ct value (33.88) as compared to the original sample (27.80). KOP cells showed no visible cytopathic effect (Table 1). All environmental surface samples were EqPPV negative.

**Table 2.** Analytic sensitivity of EqPPV qPCR.

| Copies/reaction | Ct-values of 5 parallel reactions |  |  |  |  |
| --- | --- | --- | --- | --- | --- |
| 1000 | 31.48 | 31.61 | 30.64 | 31.34 | 30.49 |
| 100 | 33.10 | 32.90 | 32.73 | 32.59 | 33.23 |
| 10 | 36.20 | 36.99 | 35.81 | 35.81 | 35.83 |
| 1 | 38.82 | No Ct | No Ct | No Ct | No Ct |
| 0 | No Ct | No Ct | No Ct | No Ct | No Ct |

### Phylogenetic analysis showed little variation between variants

EqPPV sequence was acquired from all six samples subjected to NGS (Table 1 and S1). Also, *Astroviridae*, *Parvoviridae*, and *Picobirnaviridae* were detected in two (Table S2). In total, 125 open reading frames (ORFs) were identified, and the preliminary genome draft was assembled in four fragments (Fig. 3A, Table S3). Mean amino acid (aa) identities to ORFV-SA00 and BPSV-AR02 were 67.12% and 63.84% (Table S3), whereas aa identities between EqPPV variants from this study and F14.1158H ranged 97.00–100.00%, excluding the putative chemokine-binding protein (ORF122) with aa identity of 93.09% between F14.1158H and variant A1. The nucleotide sequences of three poxvirus core genes previously identified as good candidates for phylogenetic analysis (33) - DNA polymerase (ORF025), early transcription factor VETFL (ORF083), and DNA topoisomerase type 1 (ORF062) - and envelope phospholipase (ORF011) were compared from A1, B3, F3, and J2 (Table S4). Nucleotide identities between the F14.1158H and 2021–2022 samples were between 99.20% (ORF011) and 99.48% (ORF025). Identities within the four samples varied between 99.73% (ORF011) and 100.00% (ORF083, ORF062). EqPPV variants from 2021–2022 were 74–87% identical to other PPVs and grouped together in all phylogenetic trees (Fig. S1). In addition, partial ORF093 sequence was acquired from 14 samples from nine stables by Sanger sequencing. Sequences were 100.00% identical with each other and 98.70% identical with F14.1158H (Fig. 3B).

**FIG 3.**
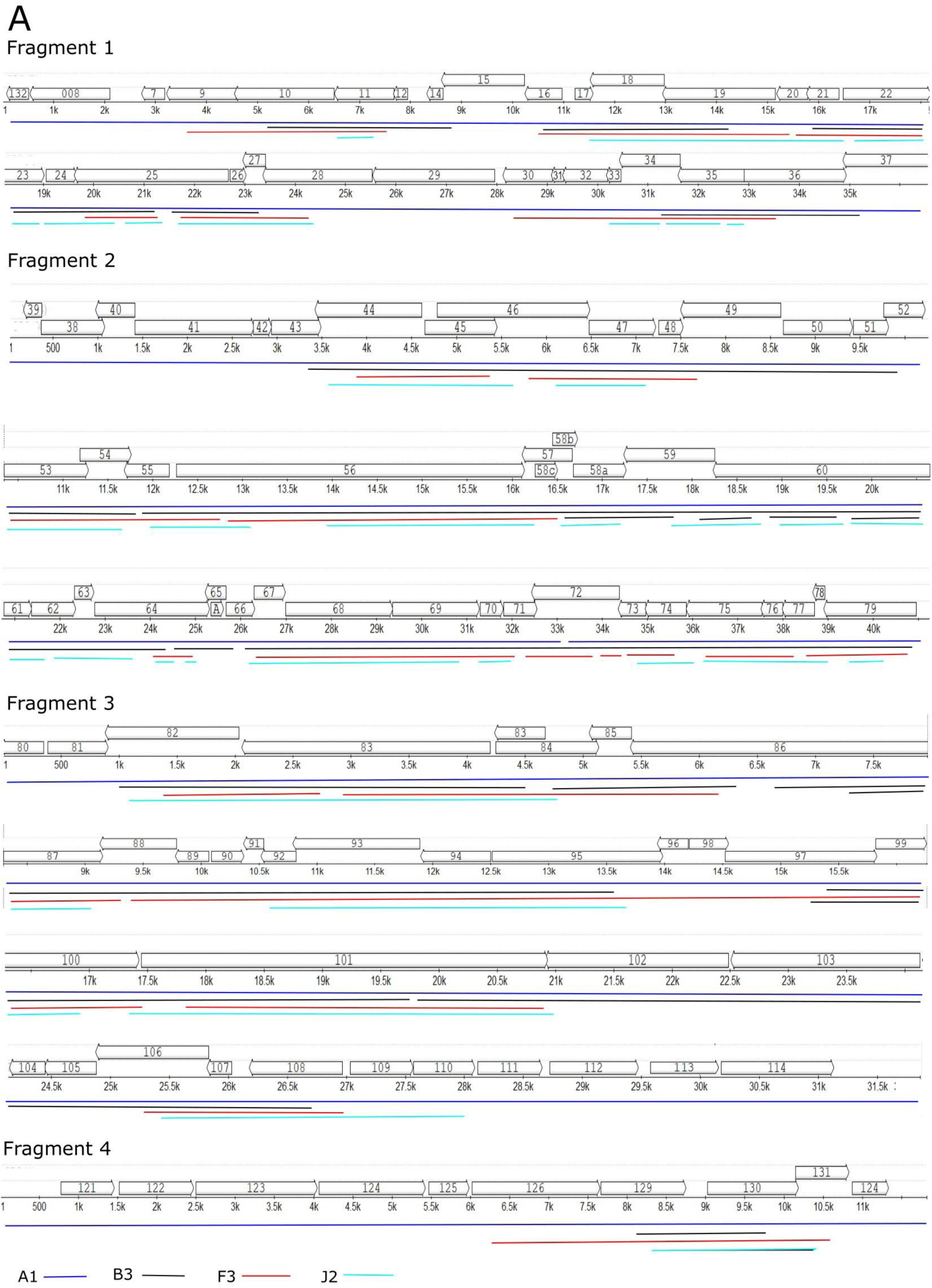
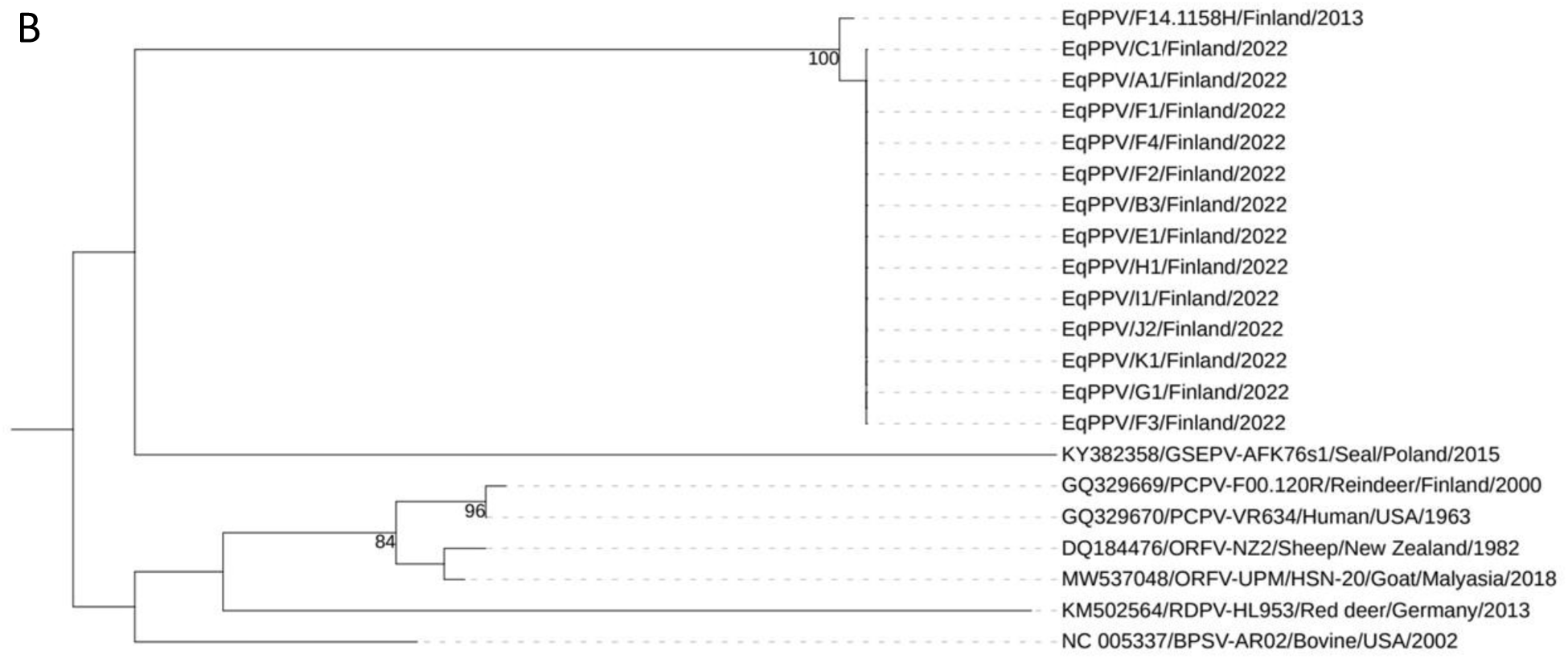
Genomic structure and phylogenetic analyses of EqPPV. A: Preliminary genome and annotation of EqPPV based on next generation sequencing results. Placement of sequence fragments of variants from four horses is indicated with coloured lines (blue: A1, black: B3, red: F3, and light blue: J2). B: Phylogenetic tree of the partial ORF093 (nt 349-872) of EqPPV variants and representatives of other PPV species. Variant labels refer to horses listed in Table 1 living at stables A-K. Tree was built with IQ-TREE 2.2.0.7 and visualized with iTOL. Boostrap values above 70 are shown next to the nodes.

### Total of 121 valid survey responses were received

Survey on epidemiology, clinical sings, treatment, distribution, and risk factors received 126 responses. Five invalid responses were discarded (one not from Finland, four outside the defined timeline). Of the remaining 121 responses, 43 (35.5%) were from affected (case) stables and 78 (64.5%) from nonaffected (control) stables (Fig. 2). Most respondents identified themselves as stable owners (93/121; 76.9%), followed by horse owners (31; 25.6%), stable personnel (14; 11.6%), trainers (11; 9.1%), veterinarians (3; 2.5%), and something else (13; 10.7%). These do not sum up to 100%, since 22.3% of the respondents had several roles. Respondents identified themselves as “horse owners” more often at the case than at the control stables (Table S5a).

### Equine pastern dermatitis was not associated with other animal species

Mean number of horses per stable was nine (median 6, range 1–50). The number did not differ between case and control stables (Table S5a). Case stables had at least one racehorse more commonly than control stables (p=0.001; Table S5a). Majority (113/121; 93.4%) of the stables had other animal species including dogs (97/121; 80.2%), cats (84/121; 69.4%), poultry (17/121; 14.0%), sheep (12/121; 9.9%), rabbits (8/121; 6.6%), cattle (3/121; 2.5%), goats (2/121; 1.7%), and swine (1/121; 0.8%). No stable had donkeys or alpacas. Of the case stables, 9.3% and 0.0%, and of the control stables, 24.4% and 7.7%, reported rats and wild ruminants (Table S5a). Differences in domestic or the mentioned wild animal species were not significant. Instead, other wild animals were reported more commonly by control stables than case stables (Table S5a).

### Practices varied as regards bedding, keeping horses outside, and hand hygiene

Using peat as bedding was more common at control stables, whereas wood-based material was more commonly used at case stables (Table S5a). Furthermore, control stables more commonly reported keeping horses outside only. Although sharing equipment between horses was more common at stables having one or more racehorse (70.7% vs. 40.9%; p=0.002), there was no difference between the case and control stables based on whether horses used individual or shared equipment (Table S5a). Use of sheep wool in leg bandages was more commonly reported by the case stables (Table S5a). Case stables also more commonly reported regularly washing or sanitizing hands between the handled horses than control stables (Table S5a).

### Quality problems in the paddock surface and attendance in race events were common

Before the onset of the clinical signs, about half of the case stables reported sandy bottom in their paddocks (Table S6a). Majority reported snow or ice cover, and about half quality problems in the paddock surface (Table S6a).

During the month before the onset of pastern dermatitis of the index case, visits of horses outside the stable were reported by 25.6%, and the arrival of a new horse, bedding lot, and feed lot by 14.0%, 7.0%, and 4.7% (Table S6a). The attendance in race events differed between the case stables (72.1%; Table S6a) and the control stables (30.8%; p<0.001), although case stables reported attendance for one month only whereas control stables for four months (December – March).

Most (25/43; 58.1%) of the respondents from the case stables believed that their horses got infected at the racetrack. Some assumed that the infection was related to farrier’s visit (4/43; 9.3%), new horse coming to the stable (4/43; 9.3%) or a horse visiting clinic or other stable (4/43; 9.3%).

### Confounding factors to be considered

Some differences detected between the case and control stables may be confounded by whether the stable kept one or more racehorse. These include keeping horses outside only, more commonly reported by control stables, but also associated with not having any racehorses (Table S5a), i.e., stables of which majority were control stables. Conversely, the use of wool bandages, proper hand hygiene, and attending race events, more commonly reported by case stables, were also associated with having one or more racehorse (Tables S5a, S6a), i.e., stables of which majority were case stables.

### One third of horses at the affected stables got pastern dermatitis manifesting as scabby, oedematous, and ulcerative skin lesions

On average, case stables had 10.3 (median 8; range 1–50) horses, of which 3.2 (median 2; range 1–20), had been affected. From 40 of the 43 case stables, median of 33.3% (mean 39.2%, range 3.8–100%) of the horses were diseased, and 3 stables (mean 5%, median 0%, 0–100%) reported still having one horse with signs. Out of 31 stables with full data concerning healing proportions, all reported diseased horses having healed and majority (mean 80.9%, median 100%, 0–100%) of them with treatment. Most often, the first clinical signs were detected in December 2021 (Fig. 4).

**FIG 4.**
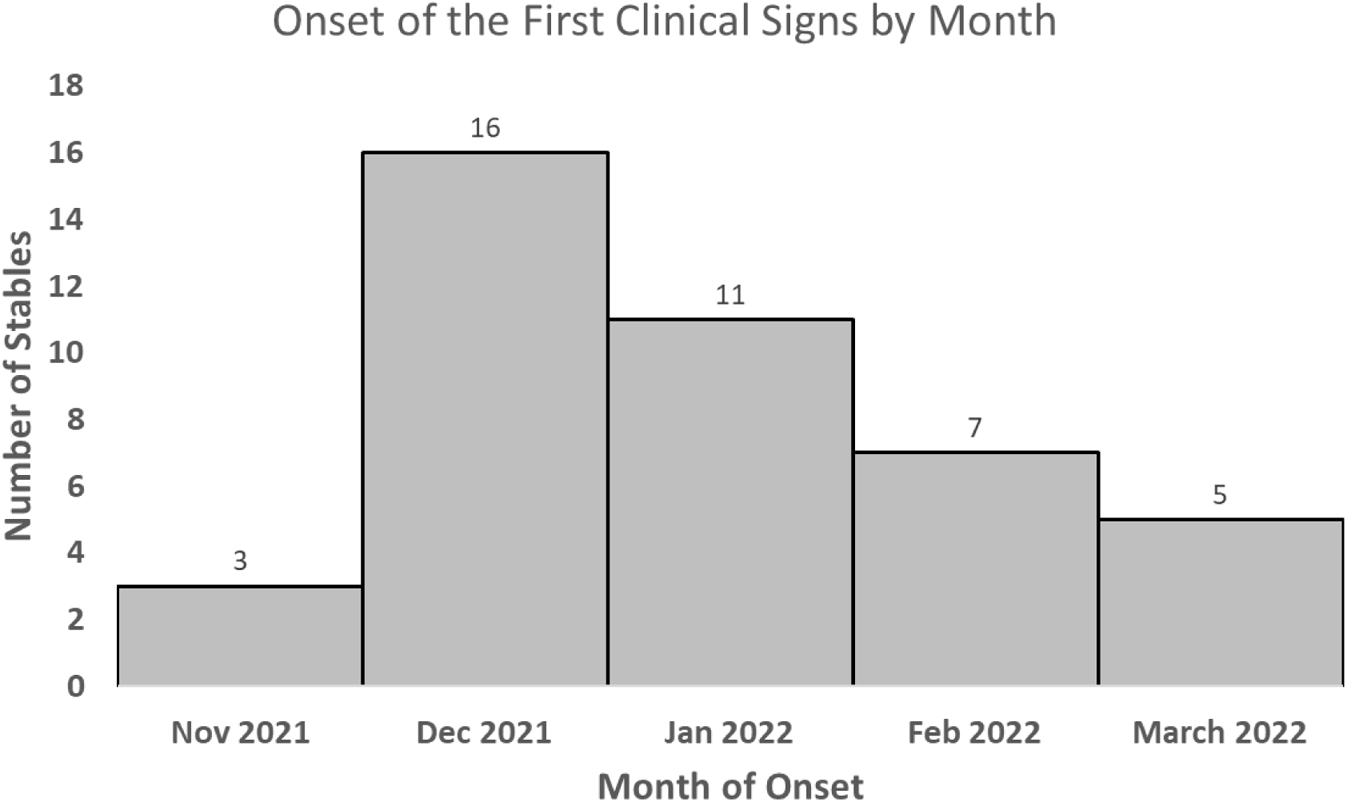
Number of case stables (N=42) reporting the onset month of pastern dermatitis in the first horse. Data were missing from one stable.

Of the 43 case stables, almost all reported scabby, oedematous, and ulcerative, and great majority excreting, erythematous, and vesicular skin lesions in the pastern region (Table S6b, Fig. 5). All these signs were reported by 41.9% of the stables (Table S6b). We did not ask about pustules, which, however, were visible in photos received (Fig. 1). There were more skin lesions outside the pastern region as well as fever and lymphangitis at case than control stables (Table S5b; p=0.06). The median duration of signs was 12–30 days (Table S6c). The median time between the onset of signs of the first and the second case was three days (Table S6c). Four (9.5%) of the 42 case stables answering reported samples being taken from the horses, but none sent to virological analyses.

**FIG 5.**
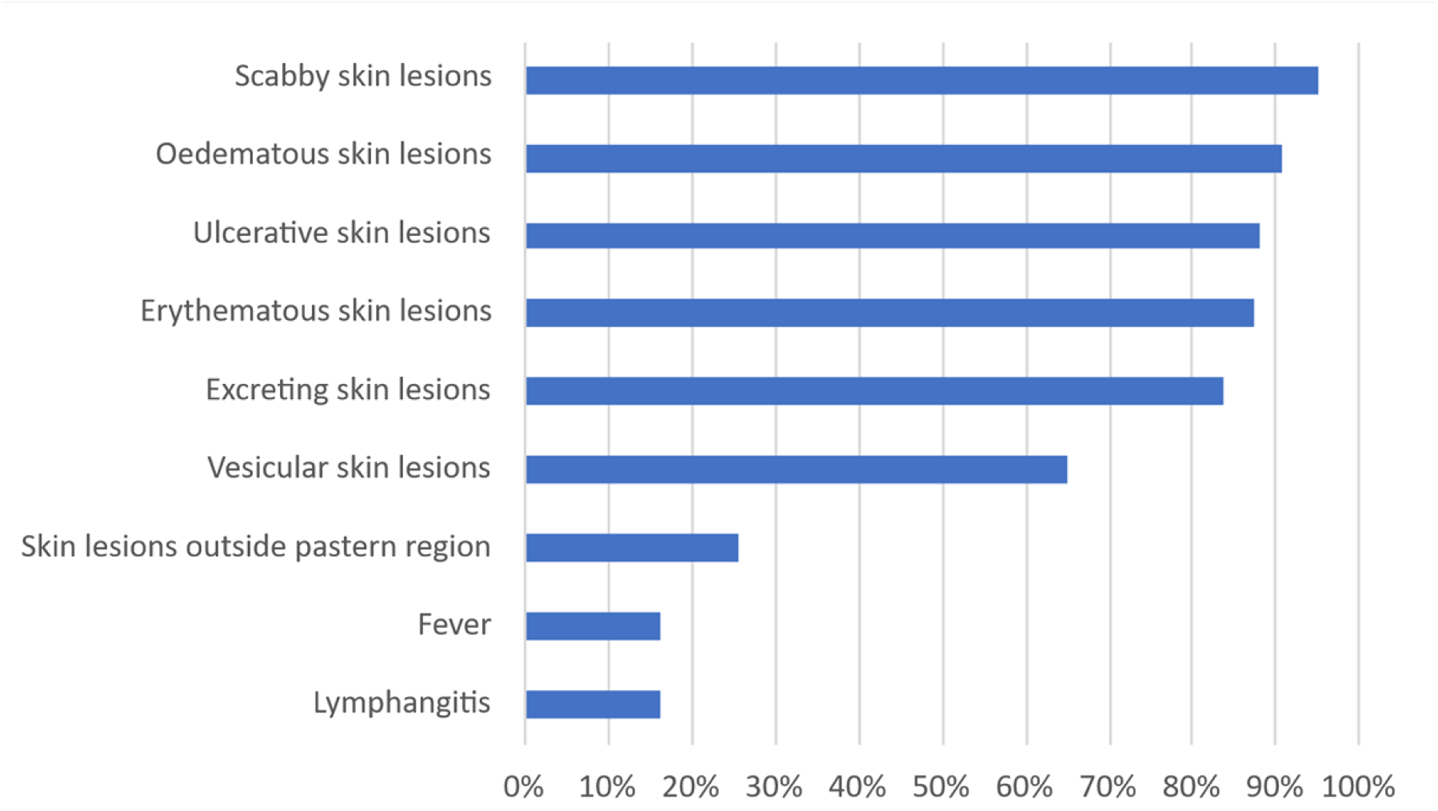
Proportion of the case stables (N=43) reporting certain clinical signs in horses. The skin lesions reported are in the pastern area unless stated otherwise.

### Long local treatment and antibiotics were needed

Clinical signs were typically treated for 10–21 days (medians for the minimum and maximum periods), varying from 0 to 210 days (Table S6c). Majority of the stables indicated local treatment sufficient for the recovery, whereas some had needed systemic antibiotic or something else (Table S6d).

Sick horses were isolated at one stable only and treated in some way by all the stables. Local topical treatment was used by most (40/43; 93.0% Fig. 6). These typically consisted of combination of three or four treatments (altogether 21 stables; 48.9%), including resin ointment, wash with antiseptic shampoo, honey ointment, wool bandages, and antibiotic ointment (Fig. 6). Fourteen stables (32.6%) gave systemic antibiotics - however, nine (20.9%) did not know if antibiotic was used (Fig. 6). Any antibiotic treatment, local or systemic, was used by more than half of the stables, whereas something else, such as non-steroidal anti-inflammatory medicines, were given at four (8.7%) (Fig. 6).

**FIG 6.**
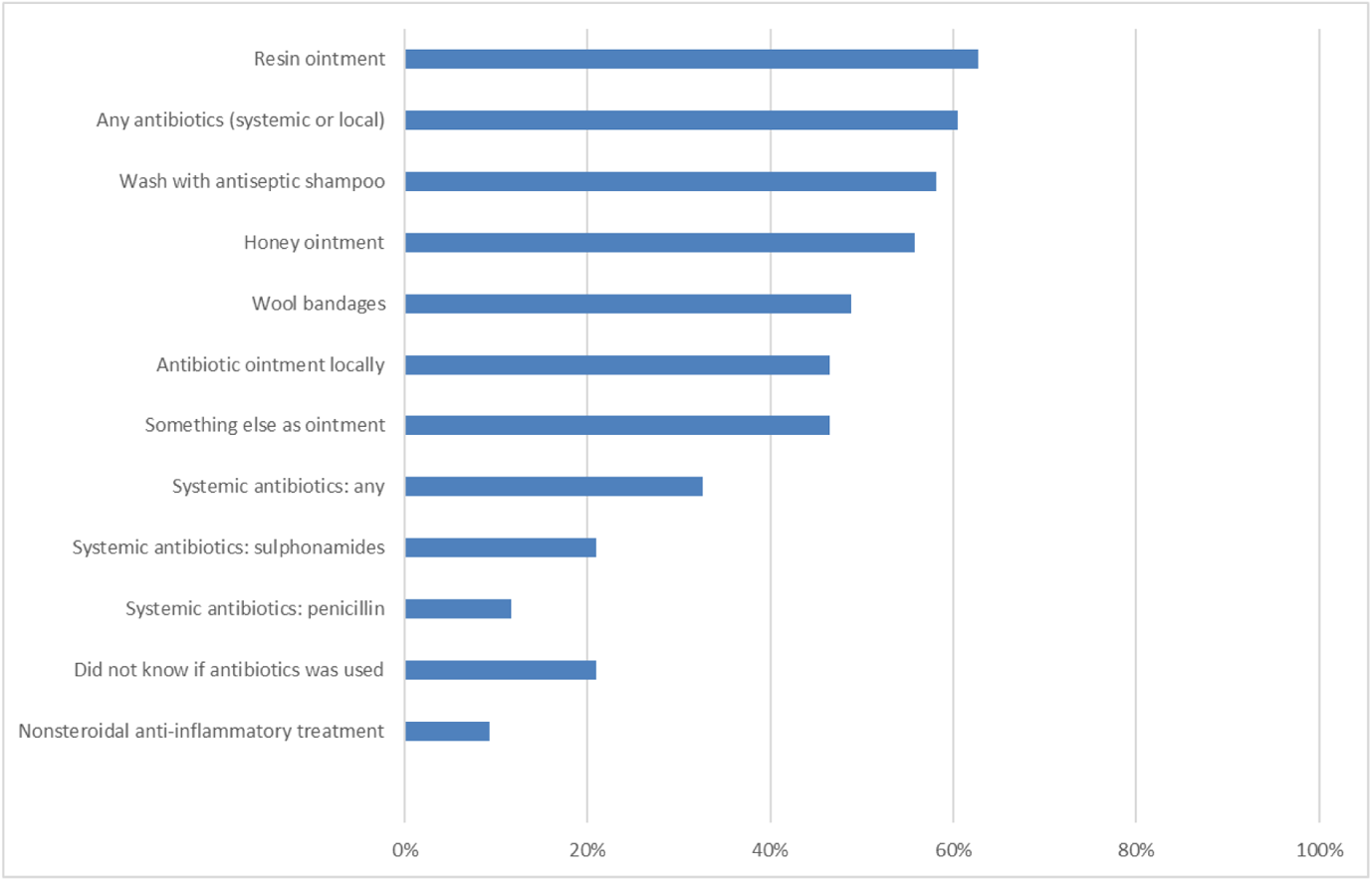
Proportion of the case stables (N=43) reporting certain treatments of horses with pastern dermatitis.

### Pastern dermatitis was associated with skin symptoms in humans

Skin lesions in animals other than horses were rare: only single cases of scabby or oedematous and four cases of erythematous skin lesions were reported by the control and none by case stables (p>0.05). Skin symptoms in humans were reported by two control (2.7%) and seven case stables (17.5%; p=0.028; Table S5c). One report of human signs from a control stable was omitted from the analysis since the free word answer revealed symptoms starting after taking care of horses at an affected neighbour stable. Erythematous and vesicular skin lesions were more commonly reported from case stables (Table S5c). All the seven case stables reported erythema, four vesicles, three swelling, two scabs or excretion, and one reported ulcer; and the control stable who detected human disease after contact with neighbouŕs diseased horses, reported all these symptoms (Fig. 7). Free-word answers reported onset of painful hand lesions 1–3 days after a glove-free contact with a horse with pastern dermatitis and recovery in 1–3 weeks.

**FIG 7.**
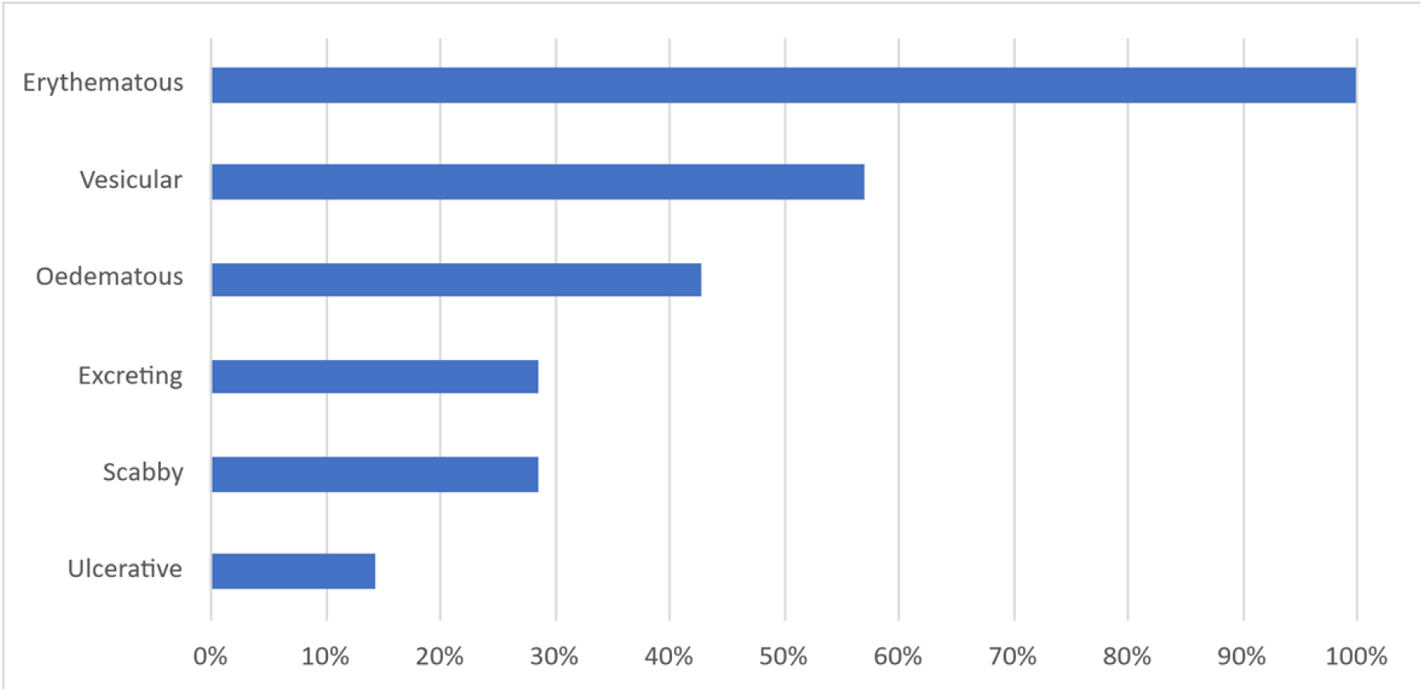
Proportion of the stables (N=8) reporting certain type of skin symptoms in humans in association with equine pastern dermatitis. Seven (case) stables had pastern dermatitis themselves and at one (control) stable, the person got skin symptoms after visiting a neighbour stable with pastern dermatitis.

## Discussion

This study delivers a detailed description of a previously undescribed outbreak of a clinically relevant equine dermatitis. The disease quickly spread all over Finland (Fig. 2, Fig. 4), causing painful lesions and prolonged welfare problems (Fig. 1). The epidemic resulted in the need of many treatments causing extra work for the personnel, and prevented training and racing, which had economic consequences to the owners. Based on the information on other PPVs and reports from stables, it also posed a zoonotic risk. We identified the likely cause of the disease, described its clinical and microbiological picture, and gained significant information about its behaviour and potential transmission routes. We also set up and validated a diagnostic PCR to enable reliable diagnostics and analysed the almost complete virus genome.

At the beginning of the outbreak, the causative agent had to be identified. The initially used pan-PPV-PCR could identify EqPPV-DNA from several but not all samples (12, 14). The more sensitive EqPPV-specific PCR later detected EqPPV in majority of the affected horses, but not in healthy controls. This combined with the fact that the same virus was detected in a sick horse in 2013 (11) and clinical signs resemble those caused by other known PPVs (Fig. 1, Fig. 5, (1, 7)) provides strong evidence that EqPPV is the causative agent of pastern dermatitis.

In addition to EqPPV, other possible causes should be considered. Our analyses did not provide any other uniform explanation for the cause than EqPPV, although, unfortunately, the skin biopsies missed the margin of lesion and healthy skin, in which changes typical for EqPPV could have been seen in histopathological analyses (11). Despite the lack of histopathological proof of PPV, it was the only uniform virus found in the NGS, which would most likely have detected other dermatopathogenic dsDNA viruses, including herpes-, papilloma- and orthopoxviruses. Bacteria are also a possible cause. However, the bacteriological results revealed no single causative species, but several pathogenic and zoonotic bacteria, such as toxin-producing *Corynebacterium diphteriae*, which were also found in a parallel study (17, submitted manuscript). The abundant growth of species originating from the normal microbiome or environment suggest secondary infection. Indeed, viral infections predispose to secondary bacterial infections, which often cause severe inflammatory changes complicating the histopathological detection of lesions caused by primary viral infection.

Full characterization of EqPPV is complicated because the virus proved challenging to isolate with monkey or ruminant cell lines previously used to isolate orthopoxvirus (26, 34) and PPVs (7). We have, however, most of the genome sequenced from clinical samples enabling building of a genome draft. The genome is missing short fragments, which is why we were unable to evaluate possibly missing genes as compared to other PPVs. Sequences from 2021–2022 indicated that virus had barely mutated. This suggests a common source of infection and is no surprise as evolutionary rate of large DNA viruses is slow (35) and previous studies have reported mutation rates of 10^-6^–10^-4^ substitutions/site/year for poxviruses (36, 37). Even though analysing the viral sequences can be used to track outbreaks and estimate transmission routes of many viruses, partial Sanger sequencing of the circulating EqPPV variants could not be applied for this outbreak due to nonexisting variation. This could, however, be tested with poxvirus genes showing more variation, e.g. ORF112.

Based on the clinical data and survey responses, pastern dermatitis in association with EqPPV seems to be oedematous and scabby pustulo-vesiculo-ulcerative skin disease localized in the pastern area (Fig.1, Table S5b, Table S6b, Fig. 5). Notably, the clinical picture is incomplete if based on the survey responses only (Fig. 5), as the respondents were mainly non-veterinarians unaware of the exact dermatological terminology. Furthermore, we missed asking about pustular or proliferative lesions in the pastern area – which should have been asked based on other data and photos.

The long duration of painful lesions led to multiple kinds of treatments, mainly local ointments (Table S6c, Table S6d, Fig. 6). Antibiotics were commonly used, which is in line with the commonness of pathogenic bacteria in the cultured samples. However, majority of the stables reported recovery without systemic antibiotics (Table S6d). We found the rare use of pain medication surprising (Fig. 6) - this welfare risk should be addressed in potential future epidemics.

Based on both the clinical information and the epidemiologic survey, this outbreak principally affected racehorses. Having racehorses and contact with racecourse were significant risk factors (Table S6a). We recognized also other potentially relevant events, actions, and visits during the month before onset, but these could not be compared with control stables since they were asked to report these for four months and we did not ask for numbers of actions, but only occurrence (Table S6a). Using wool bandages was more common at case stables and keeping horses outside only at control stables (Table S5a). However, these cannot be named as risk or protective factors because the commonness of these practices also differed between stables with/without racehorses. Usage of wool bandages may also be reactive to the outbreak. Using peat as bedding might be protective whereas wood-based bedding may be a risk factor (Table S5a). Hand hygiene practices are often suboptimal at stables (38). Better hand hygiene practices at case stables may be reactive action - we did not ask about practices used before the outbreak (Table S5a). Furthermore, proper hand hygiene was also associated with keeping racehorses. Anyhow, the importance of hygiene needs to be emphasized to prevent transmission. Surprisingly, none of the valid case stables reported sending samples to virological analyses. Therefore, the survey responses do not overlap with our laboratory analyses. However, the clinical picture (Fig. 1, Fig. 5) and geographical distribution (Fig. 2) are similar for sampled stables and responding stables.

ORFV, BPSV, and PCPV are zoonotic causing painful skin symptoms in humans. Information of GSEPV and RDPV is so scarce that zoonotic nature can’t be excluded. Here, clinical symptoms in humans were more commonly reported by case stables (Table S5c). The reported symptoms fit the clinical picture of farmyard pox and those reported from the US after equine contact in association with a novel poxvirus (Table S5c, Fig. 7; (5, 10, 39)). Unfortunately, we did not receive any human samples, so the zoonotic aspects of EqPPV remains to be verified. However, based on data from the survey and spontaneous reports, the zoonotic role of other PPVs, and the zoonotic bacteria cultured from lesions of some horses in this outbreak, a One Health approach is needed for diagnosing and biosafety measures are necessary to protect the health of people in association with equine patients.

As there is no antiviral treatment available, reliable diagnostics is even more relevant for disease control. The lack of knowledge of the causative agent and delays in diagnostics may have contributed to the efficient spread. New outbreaks will likely surface elsewhere or in Finland once the immunity decreases or animal population changes. Sensitivity of pan-PPV-PCR proved insufficient for detecting EqPPV, possibly due to four primer mismatches. Also, pan-PPV-PCR detected a substantial amount of host genome leading to only 36% of the tested dermatitis cases being confirmed EqPPV-positive whereas our new EqPPV-specific PCR was positive in 88% (23/26) of the cases. Due to the lack of reference test and variation in sampling time after the onset of signs, diagnostic sensitivity could not be reliably calculated. Based on the PCR not detecting ORFV and PCPV, it appears to be specific for EqPPV. As only two samples were tested, cross-reaction with other PPVs cannot be fully excluded. However, no other PPVs have been reported in horses and any PPV from horse samples can be considered a clinical finding.

Active participation in events may have contributed to the efficient spread of pastern dermatitis. In the future, it is important to recognize the infection early to avoid the participation of infected horses. The median interval between the first and second case at each stable (three days) indicates that direct or indirect within-stable transmission was also common. To avoid within-stable transmission, hygienic practices are necessary and more information on the modes of transmission is needed. Poxviruses can stay viable in certain conditions for prolonged periods (40), which can contribute to the spread via stable environment. Insufficient hand hygiene and biosecurity practices can contribute to the spread via humans taking care of infected horses. Other PPVs have been found from ticks and flies in infected farms (41, 42). In this outbreak, these vectors are unlikely because the outbreak occurred during Nordic winter. Based on the survey, the outbreak was not associated with any other animal species (Table S5a), but to recognize or exclude potential reservoir species, transmission routes and spill-over infections of EqPPV among wild or domestic animals, more studies and sampling are needed. The need for animal and environmental sampling and the reported clinical signs in humans associated with equine pastern dermatitis emphasize that further actions and studies should use the One Health approach.

## Acknowledgements

We thank veterinarians who participated in sampling and owners who allowed the sampling, and delivered information, and all who responded to the epidemiological survey. We thank Dr. Maria Hautaniemi, Finnish Food Authority, for the virus cultivation trial in KOP cells and her expert support when planning this study. In addition, we thank Finnish Food Authority for giving us clinical ORFV and PCPV samples for PCR optimization and Dr. Lev Levanov for the PCR positive control.

