## Supplementary for "Equine dermatitis outbreak associated with parapoxvirus"

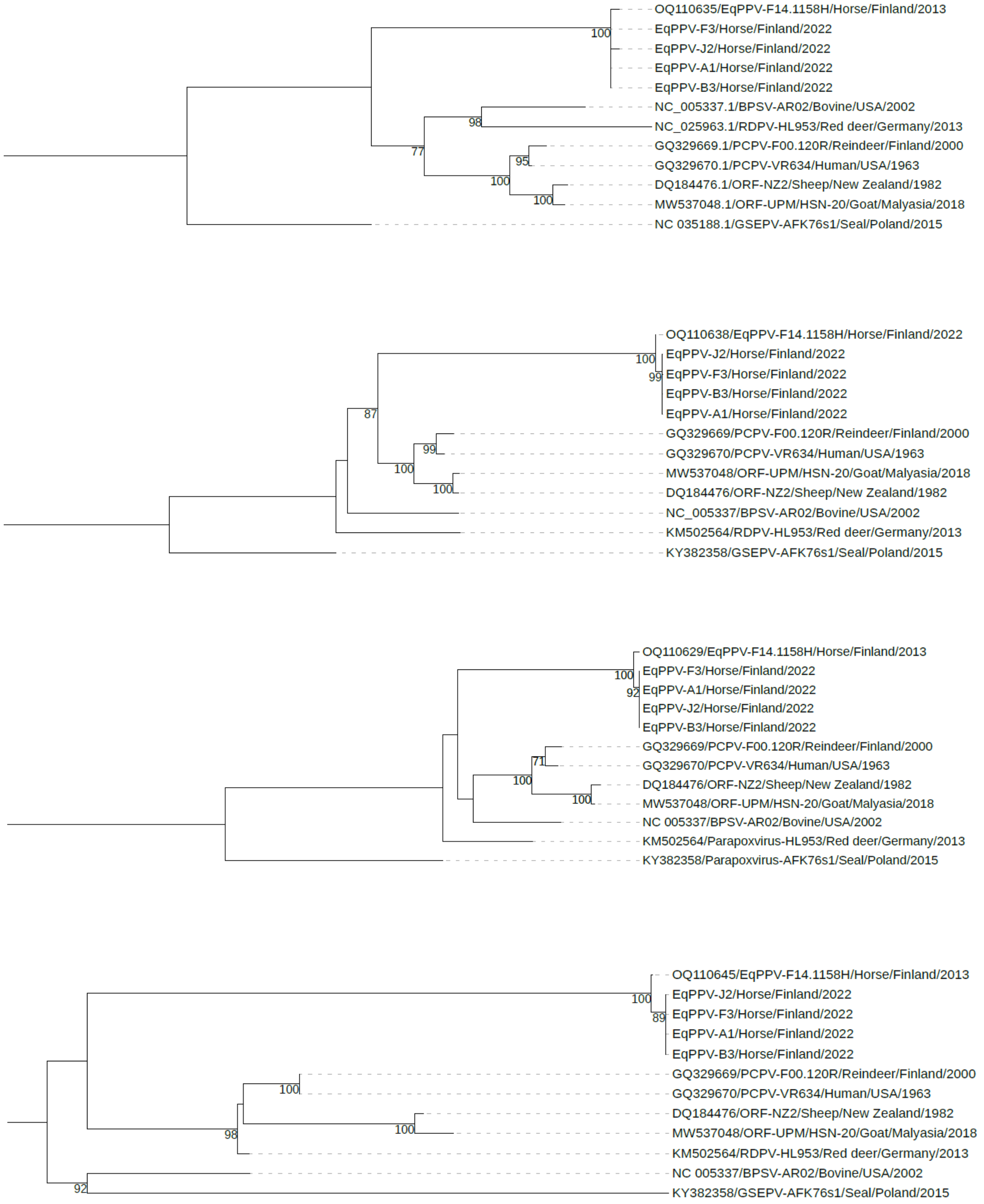


**FIG S1** Phylogenetic tree of the complete envelope phospholipase (ORF011), DNA polymerase (ORF025), early transcription factor VETFL (ORF083), and DNA topoisomerase type 1 (ORF062) genes of EqPPV strains and representatives of other PPV species. Trees were built with IQ-TREE 2.2.0.7 and visualized with iTOL. Boostrap values above 70 are shown next to the nodes.

**TABLE S1** Basic statistics from next generation sequencing

| Sample ID | Reads (N) | Reads mapping to PPVs (N) | Contigs (N) | Sequence data after assembly (kb) |
|---|---|---|---|---|
| A1 | 15430278 | 13811 | 6 | 122 |
| A2 | 22934276 | 18 | 3 | 1.3 |
| B3 | 10329094 | 3204 | 25 | 81 |
| D1 | 2080482 | 2 | 1 | 0.3 |
| F3 | 20912788 | 2256 | 47 | 76 |
| J2 | 5238710 | 1615 | 89 | 61 |

**TABLE S2** Other viruses identified in next generation sequencing

| Sample | Read (N) | Contig (N) | Lenght (kb) | Family | Genus |
|---|---|---|---|---|---|
| B3 | 89 | 6 | 3.20 | *Astroviridae* | *Mamastrovirus* |
| | 59 | 2 | 0.97 | *Parvoviridae* | *Bocaparvovirus* |
| | 13 | 3 | 1.09 | *Picobirnaviridae* | *Picobirnavirus* |
| D1 | 28 | 3 | 0.93 | *Astroviridae* | *Mamastrovirus* |
| | 10 | 1 | 0.50 | *Parvoviridae* | *Bocaparvovirus* |
| F3 | 58 | 2 | 2.04 | *Betaretrovirus* | *Retroviridae* |
| J2 | 12 | 1 | 0.46 | *Betaretrovirus* | *Retroviridae* |

**TABLE S3** PPV genes identified in EqPPV and comparison to reference strains ORFV-SA00 (AY386264) and BPSV-AR02 (NC_005337).

| ORF[1] | Predicted function | Length (nt\|aa) | Identity to ORFV-SA00 (%,)/SA00 length (aa) | Identity to BPSV-AR02 (%)/AR02 lenght (aa) | aa identity within EqPPV variants |
|---|---|---|---|---|---|
| A[2] | Virion protein-like protein | 264 \| 87 | NA | NA | NA |
| B[2] | Hypothetical protein | 522 \| 173 | NA | NA | 100 |
| ORF007 | dUTPase | 420 \| 139 | 70 (238) | 70 (163) | NA |
| ORF008 | Ankyrin repeat protein | 1536 \| 511 | 32 (530) | 32 (518) | NA |
| ORF009 | Hypothetical protein | 1323 \| 440 | 53 (452) | 54 (464) | 100 |
| ORF010 | Putative EEV maturation protein | 1920 \| 639 | 67 (643) | 66 (643) | 100 |
| ORF011 | Envelope phospholipase | 1137 \| 378 | 77–78 (378) | 75–76 (378) | 99–100 |
| ORF012 | Hypothetical protein | 249 \| 82 | 58 (89) | 43 (85) | 100 |
| ORF014 | Modified RING finger protein | 282 \| 93 | 60 (93) | 59–60 (93) | 99–100 |
| ORF015 | Hypothetical protein | 1590 \| 529 | 55 (539) | 44–45 (536) | 100 |
| ORF016 | Hypothetical protein | 693 \| 230 | 43 (259) | 44 (249) | 99 |
| ORF017 | DNA-binding phosphoprotein | 339 \| 112 | 51 (105) | 54 (105) | 100 |
| ORF018 | Poly-A polymerase catalytic subunit PAPL | 1413 \| 470 | 75 (472) | 76 (481) | 100 |
| ORF019 | Hypothetical protein | 2184 \| 727 | 63 (725) | 62 (725) | 100 |
| ORF020 | dsRNA-binding PKR inhibitor | 576 \| 191 | 64 (183) | 66 (191) | 99–100 |
| ORF021 | RNA polymerase subunit RPO30 | 609 \| 202 | 81 (193) | 76 (196) | 100 |
| ORF022 | Hypothetical protein | 1698 \| 565 | 83 (567) | 84 (566) | 100 |
| ORF023 | Membrane protein | 819 \| 272 | 75 (272) | 80 (272) | 100 |
| ORF024 | NF-kB inhibitor | 594 \| 197 | 57 (292) | 53 (227) | 100 |
| ORF025 | DNA polymerase | 3027 \| 1008 | 79 (1012) | 78 (1009) | 99–100 |
| ORF026 | Putative IMV redox protein | 306 \| 101 | 79 (96) | 80 (97) | 100 |
| ORF027 | Virion core protein | 414 \| 137 | 80 (137) | 77 (137) | 99–100 |
| ORF028 | Hypothetical protein | 2130 \| 709 | 56 (709) | 56 (700) | NA |
| ORF029 | Hypothetical protein | 2391 \| 796 | 60 (806) | 59 (807) | NA |
| ORF030 | DNA-binding virion protein | 957 \| 318 | 74 (321) | 74 (322) | 100 |
| ORF031 | Hypothetical protein | 216 \| 71 | 63 (70) | 67 (69) | 100 |
| ORF032 | DNA-binding phosphoprotein | 858 \| 285 | 60–61 (285) | 63–64 (288) | 100 |
| ORF033 | Putative IMV membrane protein | 258 \| 85 | 49 (78) | 52 (86) | 100 |
| ORF034 | Telomere-binding protein | 1170 \| 389 | 74 (389) | 71 (389) | NA |
| ORF035 | Virion core protease | 1272 \| 423 | 82 (430) | 84 (430) | 100 |
| ORF036 | RNA helicase NPH-II | 2013 \| 670 | 73 (683) | 71 (684) | 100 |
| ORF037 | Putative metalloprotease | 1794 \| 597 | 73 (603) | 70 (603) | NA |
| ORF038 | Late transcription elongation factor | 702 \| 233 | 59 (231) | 63 (233) | 100 |
| ORF039 | Hypothetical protein | 183 \| 60 | 47 (110) | 50 (111) | 100 |

| ORF | Predicted function | Length (nt\|aa) | Identity to ORFV-SA00 (%,)/SA00 length (aa) | Identity to BPSV-AR02 (%)/AR02 lenght (aa) | aa identity within EqPPV variants |
|---|---|---|---|---|---|
| ORF040 | Putative glutaredoxin 2 | 420 \| 139 | 69 (137) | 72 (137) | 100 |
| ORF041 | Hypothetical protein | 1320 \| 439 | 65–66 (452) | 63 (441) | 97–100 |
| ORF042 | RNA polymerase subunit RPO7 | 192 \| 63 | 79 (63) | 75 (63) | 100 |
| ORF043 | Hypothetical protein | 546 \| 181 | 62 (185) | 61 (192) | 100 |
| ORF044 | Virion core protein | 1173 \| 390 | 59–58 (398) | 58–57 (391) | 100 |
| ORF045 | ORF045 late transcription factor VLTF-1 | 801 \| 266 | 93 (266) | 94 (266) | 100 |
| ORF046 | Myristylated protein | 1692 \| 563 | 67 (344) | 73 (333) | 100 |
| ORF047 | Putative myristylated IMV envelope protein | 735 \| 244 | 79 (244) | 78–79 (244) | 100 |
| ORF048 | Hypothetical protein | 267 \| 88 | 59–60 (90) | 60 (91) | 99 |
| ORF049 | Hypothetical protein | 1095 \| 364 | 62 (376) | 60 (417) | 100 |
| ORF050 | DNA-binding virion core protein VP8 | 762 \| 253 | 73 (256) | 71 (259) | 100 |
| ORF051 | Putative membrane protein | 384 \| 127 | 59 (129) | 66 (128) | 100 |
| ORF052 | Putative IMV membrane protein | 453 \| 150 | 73 (151) | 69 (151) | NA |
| ORF053 | Poly-A polymerase small subunit VP39 | 1014 \| 337 | 79 (336) | 81 (337) | 100 |
| ORF054 | RNA polymerase subunit RPO22 | 561 \| 186 | 84 (186) | 78 (186) | 99–100 |
| ORF055 | Late membrane protein | 477 \| 158 | 79–80 (167) | 76–77 (167) | 99–100 |
| ORF056 | RNA polymerase subunit RPO147 | 3867 \| 1288 | 86–87 (1289) | 87 (1289) | 100 |
| ORF057 | Putative protein-tyrosine phosphatase | 540 \| 179 | 64–65 (181) | 67–68 (179) | 99–100 |
| ORF058a[3] | Hypothetical protein | 576 \| 191 | 81 (321) | 82 (194) | 100 |
| ORF058b[3] | Hypothetical protein | 267 \| 88 | 60 (321) | NA | NA |
| ORF058c[3] | Hypothetical protein | 189 \| 62 | 73 (321) | NA | NA |
| ORF059 | Putative IMV protein VP55 | 993 \| 330 | 65 (342) | 63–64 (340) | 97–100 |
| ORF060 | RNA polymerase-associated protein RAP94 | 2397 \| 798 | 80 (804) | 80 (803) | 100 |
| ORF061 | Late transcription factor VLTF-4 | 603 \| 200 | 52 (227) | 59 (242) | 100 |
| ORF062 | DNA topoisomerase type I | 963 \| 320 | 77–78 (318) | 77 (320) | 99–100 |
| ORF063 | Hypothetical protein | 414 \| 137 | 58 (138) | 56 (138) | 100 |
| ORF064 | mRNA capping enzyme large subunit | 2523 \| 840 | 80 (841) | 80–81 (841) | 99–100 |
| ORF065 | Virion protein | 426 \| 141 | 67 (156) | 68 (158) | 100 |
| ORF066 | Virion protein | 630 \| 209 | 54 (291) | 56 (225) | 100 |
| ORF067 | Uracil DNA glycosidase | 678 \| 225 | 83–84 (231) | 84 (247) | 100 |
| ORF068 | NTPase | 2361 \| 786 | 84 (787) | 84 (788) | 100 |

| ORF | Predicted function | Length (nt\|aa) | Identity to ORFV-SA00 (%,)/SA00 length (aa) | Identity to BPSV-AR02 (%)/AR02 lenght (aa) | aa identity within EqPPV variants |
|---|---|---|---|---|---|
| ORF069 | Early transcription factor VETFs | 1911 \| 636 | 89 (635) | 89–90 (650) | 100 |
| ORF070 | RNA polymerase subunit RPO18 | 489 \| 162 | 80 (190) | 83 (176) | 100 |
| ORF071 | NPH-PPH downregulator | 720 \| 239 | 72 (224) | 68 (223) | 100 |
| ORF072 | Transcription termination factor NPH-I | 1098 \| 635 | 81 (638) | 80 (638) | 100 |
| ORF073 | Hypothetical protein | 570 \| 189 | 60–61 (188) | 59–60 (194) | 99–100 |
| ORF074 | mRNA capping enzyme small subunit | 873 \| 290 | 84–85 (300) | 85–86 (290) | 100 |
| ORF075 | Putative rifampicin resistance protein | 1638 \| 545 | 84 (545) | 86 (545) | 100 |
| ORF076 | Late transcription factor VLTF-2 | 447 \| 148 | 80 (150) | 78 (150) | 100 |
| ORF077 | Late transcription factor VLTF-3 | 675 \| 224 | 94 (224) | 93 (224) | 100 |
| ORF078 | Thioredoxin-like protein | 237 \| 78 | 71 (83) | 71 (80) | 100 |
| ORF079 | Virion core protein P4b precursor | 2007 \| 668 | 69 (675) | 72 (683) | 100 |
| ORF081 | RNA polymerase subunit RPO19 | 513 \| 170 | 64 (173) | 68 (171) | 99–100 |
| ORF082 | Hypothetical protein | 1137 \| 378 | 71 (378) | 70–71 (384) | 99–100 |
| ORF083a[3] | Early transcription factor 82 kDa subunit | 2124 \| 707 | 84 (841) | 84 (706) | 100 |
| ORF083b[3] | Early transcription factor VETFL | 420 \| 139 | 80 (841) | NA | 100 |
| ORF084 | Intermediate transcription factor VITF-3 | 879 \| 292 | 80 (303) | 81 (307) | 100 |
| ORF085 | Late virion membrane protein | 348 \| 115 | 73 (93) | 75 (95) | 100 |
| ORF086 | Virion core protein P4a precursor | 2718 \| 905 | 70 (905) | 72 (908) | 100 |
| ORF087 | Hypothetical protein | 996 \| 331 | 85 (336) | 83–84 (345) | 100 |
| ORF088 | Virion core protein | 645 \| 214 | 53–54 (269) | 54 (223) | 100 |
| ORF089 | Virion membrane protein | 276 \| 91 | 53 (92) | 49–51 (78) | 99–100 |
| ORF090 | Putative IMV phosphorylated membrane | 276 \| 91 | 68 (91) | 69 (90) | 100 |
| ORF091 | Putative IMV membrane protein | 162 \| 53 | 68 (53) | 75 (53) | 100 |
| ORF092 | Hypothetical protein | 288 \| 95 | 54 (89) | 57–58 (92) | 97–100 |

| ORF | Predicted function | Length (nt\|aa) | Identity to ORFV-SA00 (%,)/SA00 length (aa) | Identity to BPSV-AR02 (%)/AR02 lenght (aa) | aa identity within EqPPV variants |
|---|---|---|---|---|---|
| ORF093 | Myristylated protein | 1080 \| 359 | 75 (358) | 74 (359) | 100 |
| ORF094 | Phosphorylated IMV membrane protein | 591 \| 196 | 73–74 (196) | 76 (201) | 99–100 |
| ORF095 | DNA helicase | 1464 \| 487 | 79 (488) | 80–81 (489) | 99–100 |
| ORF096 | Putative Zn-finger protein | 252 \| 83 | 59 (90) | 63 (83) | 100 |
| ORF097 | DNA polymerase processivity factor | 1302 \| 433 | 65 (429) | 64 (426) | 100 |
| ORF098 | Hypothetical protein | 330 \| 109 | 66 (108) | 74 (164) | 100 |
| ORF099 | Holliday junction resolvase | 435 \| 144 | 88 (146) | 89 (146) | 100 |
| ORF100 | Intermediate transcription factor VITF-3 | 1146 \| 381 | 78 (380) | 80 (381) | 100 |
| ORF101 | DNA-directed RNA polymerase 132 kDa polypeptide | 3480 \| 1159 | 90 (1160) | 90 (1161) | 100 |
| ORF102 | A type inclusion protein | 1554 \| 517 | 72 (520) | 72 (520) | 99–100 |
| ORF103 | A type inclusion protein | 1608 \| 535 | 61 (616) | 62 (519) | 97–100 |
| ORF104 | Fusion protein | 288 \| 95 | 58 (90) | 55 (89) | 98–100 |
| ORF105 | Hypothetical protein | 423 \| 140 | 79 (140) | 75 (140) | 100 |
| ORF106 | RNA polymerase subunit RPO35 | 942 \| 313 | 66 (314) | 68 (319) | 99–100 |
| ORF107 | Virion morhogenesis | 195 \| 64 | 60 (60) | 69 (61) | 100 |
| ORF108 | DNA packaging protein/ATPase | 774 \| 257 | 88 (274) | 89 (259) | 100 |
| ORF109 | EEV glycoprotein | 525 \| 174 | 48 (164) | 51 (162) | 100 |
| ORF110 | EEV glycoprotein | 513 \| 170 | 48 (167) | 41 (167) | 99–100 |
| ORF111 | Hypothetical protein | 543 \| 180 | 54 (179) | 54 (184) | 99–100 |
| ORF112 | Chemocine-binding protein | 744 \| 247 | 28 (288) | 31 (297) | 93–100 |
| ORF113 | Hypothetical protein | 522 \| 173 | 45 (100) | 32 (199) | 100 |
| ORF114 | Hypothetical protein | 948 \| 315 | 52 (344) | 46 (331) | 99–100 |
| ORF121 | NF-kappa pathway inhibitor | 702 \| 233 | 32 (302) | 42 (269) | 99–100 |
| ORF122 | Hypothetical protein | 942 \| 313 | 48 (323) | 50 (322) | 100 |
| ORF123 | Ankyrin repeat protein | 1536 \| 511 | 53 (525) | 59 (517) | 100 |
| ORF124a | Hypothetical protein | 1347 \| 448 | 47 (532) | 48 (506) | NA |
| ORF125 | Hypothetical protein | 501 \| 166 | 48 (173) | 40 (177) | NA |
| ORF126 | Ankyrin repeat protein | 1620 \| 539 | 48 (173) | 40 (177) | NA |
| ORF129 | Ankyrin repeat protein | 1077 \| 358 | 46 (497) | 44 (506) | NA |
| ORF130 | Putative serine/threonine protein kinase | 1146 \| 381 | 47 (516) | 48 (515) | 100 |
| ORF131 | Membrane protein | 675 \| 224 | 83 (498) | 83 (497) | 100 |
| ORF132 | Vascular endothelial growth factor-like protein | 405 \| 134 | 62 (226) | 61 (224) | NA |
| Mean amino acid similarity (%)/standard deviation | | | 67.12/13.98 | 63.84/15.98 | |

[1] According to Delhon et al. (2004, 10.1128/jvi.78.1.168-177.2004)

[2] Gene was not annotated by Delhon et al. (2004, 10.1128/jvi.78.1.168-177.2004)

[3] Gene appeared in two or more fragments as compared to reference strain

**TABLE S4** P-distances between EqPPV variants and representatives of other PPVs

| | GQ329669/PCPV-F00.120R | GQ329670/PCPV-VR634 | MW537048/ORF-UPM/HSN-20 | DQ184476/ORF-NZ2/ | NC_005337/BPSV-AR02 | KM502564/RDPV-HL953 | KY382358/GSEPV-AFK76s1 | EqPPV-F14.1158 | EqPPV-A1 | EqPPV-B3 | EqPPV-F3 |
|---|---|---|---|---|---|---|---|---|---|---|---|
| **DNA polymerase (ORF025)** | | | | | | | | | | | |
| GQ329670/PCPV-VR634 | 0.02 | | | | | | | | | | |
| MW537048/ORF-UPM/HSN-20 | 0.06 | 0.05 | | | | | | | | | |
| DQ184476/ORF-NZ2 | 0.06 | 0.05 | 0.01 | | | | | | | | |
| NC_005337/BPSV-AR02 | 0.13 | 0.12 | 0.13 | 0.13 | | | | | | | |
| KM502564/RDPV-HL953 | 0.13 | 0.12 | 0.13 | 0.13 | 0.14 | | | | | | |
| KY382358/GSEPV-AFK76s1 | 0.21 | 0.20 | 0.21 | 0.21 | 0.21 | 0.21 | | | | | |
| EqPPV-F14.1158H | 0.18 | 0.17 | 0.18 | 0.18 | 0.19 | 0.19 | 0.26 | | | | |
| EqPPV-A1 | 0.18 | 0.17 | 0.18 | 0.18 | 0.19 | 0.19 | 0.26 | 0.01 | | | |
| EqPPV-B3 | 0.17 | 0.16 | 0.17 | 0.17 | 0.18 | 0.19 | 0.25 | 0.00 | 0.00 | | |
| EqPPV-F3 | 0.18 | 0.17 | 0.18 | 0.18 | 0.19 | 0.20 | 0.26 | 0.01 | 0.00 | 0.00 | |
| EqPPV-J2 | 0.17 | 0.16 | 0.17 | 0.16 | 0.18 | 0.18 | 0.24 | 0.00 | 0.00 | 0.00 | 0.00 |
| **Viral envelope phospholipase (ORF011)** | | | | | | | | | | | |
| GQ329670/PCPV-VR634 | 0.01 | | | | | | | | | | |
| MW537048/ORF-UPM/HSN-20 | 0.06 | 0.05 | | | | | | | | | |
| DQ184476/ORF-NZ2 | 0.06 | 0.05 | 0.02 | | | | | | | | |
| NC_005337/BPSV-AR02 | 0.15 | 0.14 | 0.15 | 0.15 | | | | | | | |
| KM502564/RDPV-HL953 | 0.17 | 0.16 | 0.17 | 0.17 | 0.14 | | | | | | |
| KY382358/GSEPV-AFK76s1 | 0.22 | 0.21 | 0.22 | 0.22 | 0.22 | 0.24 | | | | | |
| EqPPV-F14.1158H | 0.19 | 0.18 | 0.19 | 0.19 | 0.19 | 0.22 | 0.24 | | | | |
| EqPPV-A1 | 0.19 | 0.18 | 0.19 | 0.18 | 0.19 | 0.22 | 0.24 | 0.01 | | | |
| EqPPV-B3 | 0.19 | 0.18 | 0.19 | 0.19 | 0.18 | 0.21 | 0.23 | 0.01 | 0.00 | | |
| EqPPV-F3 | 0.19 | 0.18 | 0.19 | 0.18 | 0.19 | 0.22 | 0.24 | 0.01 | 0.00 | 0.00 | |
| EqPPV-J2 | 0.18 | 0.17 | 0.18 | 0.17 | 0.17 | 0.19 | 0.20 | 0.01 | 0.01 | 0.01 | 0.01 |
| **Early transcription factor VETFL (ORF083)** | | | | | | | | | | | |
| GQ329670/PCPV-VR634 | 0.02 | | | | | | | | | | |
| MW537048/ORF-UPM/HSN-20 | 0.05 | 0.05 | | | | | | | | | |
| DQ184476/ORF-NZ2 | 0.05 | 0.05 | 0.01 | | | | | | | | |
| NC_005337/BPSV-AR02 | 0.09 | 0.09 | 0.11 | 0.11 | | | | | | | |
| KM502564/RDPV-HL953 | 0.10 | 0.10 | 0.12 | 0.12 | 0.10 | | | | | | |
| KY382358/GSEPV-AFK76s1 | 0.20 | 0.20 | 0.21 | 0.21 | 0.20 | 0.20 | | | | | |
| EqPPV-F14.1158H | 0.13 | 0.13 | 0.14 | 0.14 | 0.13 | 0.13 | 0.22 | | | | |
| EqPPV-A1 | 0.13 | 0.13 | 0.14 | 0.14 | 0.13 | 0.13 | 0.22 | 0.01 | | | |
| EqPPV-B3 | 0.13 | 0.13 | 0.14 | 0.14 | 0.13 | 0.13 | 0.22 | 0.01 | 0.00 | | |
| EqPPV-F3 | 0.14 | 0.13 | 0.14 | 0.14 | 0.13 | 0.14 | 0.22 | 0.01 | 0.00 | 0.00 | |
| EqPPV-J2 | 0.13 | 0.13 | 0.14 | 0.14 | 0.13 | 0.13 | 0.22 | 0.01 | 0.00 | 0.00 | 0.00 |

| | DNA topoisomerase type 1 (ORF062) | | | | | | | | | | |
|---|---|---|---|---|---|---|---|---|---|---|---|
| GQ329670/PCPV-VR634 | 0.00 | | | | | | | | | | |
| MW537048/ORF-UPM/HSN-20 | 0.08 | 0.08 | | | | | | | | | |
| DQ184476/ORF-NZ2 | 0.09 | 0.09 | 0.02 | | | | | | | | |
| NC_005337/BPSV-AR02 | 0.14 | 0.14 | 0.15 | 0.16 | | | | | | | |
| KM502564/RDPV-HL953 | 0.03 | 0.03 | 0.07 | 0.07 | 0.12 | | | | | | |
| KY382358/GSEPV-AFK76s1 | 0.21 | 0.21 | 0.21 | 0.21 | 0.19 | 0.21 | | | | | |
| EqPPV-F14.1158H | 0.20 | 0.20 | 0.19 | 0.19 | 0.20 | 0.18 | 0.25 | | | | |
| EqPPV-A1 | 0.20 | 0.20 | 0.19 | 0.19 | 0.20 | 0.18 | 0.25 | 0.01 | | | |
| EqPPV-B3 | 0.20 | 0.20 | 0.19 | 0.19 | 0.20 | 0.18 | 0.25 | 0.01 | 0.00 | | |
| EqPPV-F3 | 0.20 | 0.20 | 0.19 | 0.19 | 0.20 | 0.18 | 0.25 | 0.01 | 0.00 | 0.00 | |
| EqPPV-J2 | 0.20 | 0.20 | 0.19 | 0.19 | 0.20 | 0.18 | 0.25 | 0.01 | 0.00 | 0.00 | 0.00 |

**TABLE S5** Comparisons between stables with pastern dermatitis signs (case) and without pastern dermatitis signs (control)

**TABLE S5a** Comparison between stables with pastern dermatitis signs (cases, N=43, 35.5%) and without signs (controls, N=78, 64.5%): horses, respondents[1], wild animals, and practices in the stable

| Factors in analysis | Case stables | | Control stables | | p-value[2] | FDR-corrected p-value |
|---|---|---|---|---|---|---|
| | n[3] | mean, median, range | n | mean, median, range | | |
| Number of horses | 43 | 10.3, 8, 1–50 | 77[4] | 9.9, 6, 1–40 | 0.179[5] | 0.203 |
| | | % (95% CI[6]) | | % (95% CI) | | |
| Respondent a horse owner | 17 | 39.5 (26.4–54.4) | 14 | 17.9 (11.0–27.9) | 0.016 | 0.034 |
| Respondent a stable owner | 28 | 65.1 (50.1–77.6) | 65 | 83.3 (73.5–90.0) | 0.041 | 0.077 |
| Respondent stable personnel | 8 | 18.6 (9.7–32.6) | 6 | 7.7 (3.6–15.8) | 0.083 | 0.107 |
| Respondent a trainer | 7 | 16.3 (8.1–30.0) | 4 | 5.1 (2.0– 12.5) | 0.052 | 0.082 |
| Respondent attending veterinarian | 2 | 4.7 (1.3–15.5) | 1 | 1.3 (0.2–6.9) | 0.287 | 0.305 |
| Respondent something else | 8 | 18.6 (9.7–32.6) | 5 | 6.4 (2.8–14.1) | 0.062 | 0.088 |
| 1 or more racehorses | 38 | 88.4 (75.5–94.9) | 37 | 47.4 (36.7–58.4) | <0.001 | 0.001 |
| Contact with wild rats | 4 | 9.3 (3.7–21.6) | 19 | 24.4 (16.2–34.9) | 0.053 | 0.082 |
| Contact with wild ruminants[7] | 0 | 0 (0–8.2) | 6 | 7.7 (3.6–15.8) | 0.088 | 0.107 |
| Contact with other wild animals[8] | 1 | 2.3 (0.4–12.1) | 16 | 20.5 (13.0–30.8) | 0.005 | 0.012 |
| Use of peat as bedding material[9] | 5 | 11.6 (5.1–24.5) | 38 | 48.7 (38.0–59.6) | <0.001 | 0.001 |
| Use of wood-based bedding material | 39 | 90.7 (78.4–96.3) | 51 | 65.4 (54.3–75.0) | 0.002 | 0.009 |
| Kept only outside[10] | 1 | 2.3 (0.4–12.1) | 16 | 20.5 (13.0–30.8) | 0.005 | 0.012 |
| Sharing equipment | 28 | 65.1 (50.1–77.6) | 43[11] | 56.6 (45.4–67.1) | 0.438 | 0.438 |
| Use of sheep wool in leg bandages[12] | 27 | 62.8 (47.9–75.6) | 24[13] | 31.2 (21.9–42.2) | 0.001 | 0.006 |
| Washing or sanitizing hands between horses[14] | 30[15] | 71.4 (56.4–82.8) | 34 | 43.6 (33.1–54.6) | 0.004 | 0.012 |

[1] The proportions of different respondent types do not sum up to 100%, since as many as 27 (22.3%) respondents had more than one role.
[2] Fisher's exact test p-value, unless otherwise indicated
[3] Numbers of stables out of all stables in cases or controls in this analysis
[4] One farm did not give this information
[5] Independent samples Mann-Whitney U test p-value
[6] Confidence interval
[7] Wild ruminants: deer or moose
[8] Such as wild rabbit, fox, raccoon dog, wolverine, wolf, mice, geese, cranes, jackdaws and other birds (other than rats and wild ruminants)
[9] Peat used alone or in addition to other bedding material
[10] Keeping horses only outside was also significantly more common in stables without racehorses (p<0.001) than in other stables
[11] Only 76 answered
[12] Use of sheep wool in leg bandages was also significantly more common in stables having 1 or more racehorses (p<0.001) than in other stables; however, we did not ask if the case stables started to use these after the outbreak.
[13] only 77 answered
[14] Whatever washing or disinfection (or all have their own caretakers) compared to others; this factor was also significantly more common in stables having 1 or more racehorses (p=0.014) than in other stables; however, we did not ask if the case stables started this practice after the outbreak.
[15] Only 42 answered

**TABLE S5b** Comparison of signs in horses elsewhere than in the pastern region between stables with (cases) and without (control) pastern dermatitis signs

| Sign | Case stables (N=43, 35.5%) | | Control stables (N=78, 64.5%) | | p-value[3] | FDR-corrected p-value |
|---|---|---|---|---|---|---|
| | n[1] | % (95% CI[2]) | n | % (95% CI) | | |
| Any kind of skin lesion outside the pastern region | 11 | 25.6 (14.9–40.2) | 6 | 7.7 (3.6–15.8) | 0.012 | 0.060 |
| Ulcers | 3 | 7.0 (2.0–18.6) | 0 | 0 (0–4.7) | 0.043 | 0.061 |
| Vesicles | 3 | 7.0 (2.0–18.6) | 0 | 0 (0–4.7) | 0.043 | 0.061 |
| Excreting lesions | 3 | 7.0 (2.0–18.6) | 0 | 0 (0–4.7) | 0.043 | 0.061 |
| Oedematous lesions | 3 | 7.0 (2.0–18.6) | 0 | 0 (0–4.7) | 0.043 | 0.061 |
| Proliferative lesions | 2 | 4.7 (1.3–15.5) | 0 | 0 (0–4.7) | 0.124 | 0.155 |
| Scabby lesions | 6 | 14.0 (6.6–27.3) | 6 | 7.7 (3.6–15.8) | 0.343 | 0.381 |
| Fever | 7 | 16.3 (8.1–30.0) | 2 | 2.6 (0.7–8.9) | 0.010 | 0.060 |
| Lymphangitis | 7 | 16.3 (8.1–30.0) | 3 | 3.8 (1.3–10.7) | 0.033 | 0.061 |
| Lameness | 6 | 14.0 (6.6–27.3) | 7 | 9.0 (4.4–17.4) | 0.541 | 0.541 |

[1] Numbers of stables out of all stables in cases or controls in this analysis
[2] Confidence interval
[3] Fisher's exact test p-value

**TABLE S5c** Comparison of symptoms in humans between stables with pastern dermatitis signs (cases) and without signs (controls)

| Symptom | Case stables (N=43, 35.8%) | | Control stables (N=77[1], 64.2%) | | p-value[4] | FDR-corrected p-value |
|---|---|---|---|---|---|---|
| | n/N[2] | % (95% CI[3]) | n/N | % (95% CI) | | |
| Ulcers | 1/40 | 2.5 (0.4–12.9) | 2/75 | 2.7 (0.7–9.2) | 1.000 | 1.000 |
| Vesicles | 4/40 | 10.0 (4.0–23.1) | 0/74 | 0.0 (0.0–4.9) | 0.014 | 0.033 |
| Excreting lesions | 2/40 | 5.0 (1.4–16.5) | 0/75 | 0 (0–4.9) | 0.119 | 0.139 |
| Oedematous lesions | 3/40 | 7.5 (2.6–19.9) | 0/75 | 0 (0–4.9) | 0.040 | 0.070 |
| Scabby lesions | 2/39 | 5.1 (1.4–16.9) | 0/75 | 0 (0–4.9) | 0.115 | 0.139 |
| Erythematous skin lesions | 7/38 | 18.4 (9.2–33.4) | 1/75 | 1.3 (0.2–7.2) | 0.002 | 0.014 |
| Any kind of skin lesion | 7/40 | 17.5 (8.8–32.0) | 2/75 | 2.7 (0.7–9.2) | 0.008 | 0.028 |

[1] One control stable was omitted from this analysis, because the respondent informed that human symptoms seen at their stable were caught from a neighbour stable affected with pastern dermatitis
[2] Numbers of stables out of all stables in cases or controls in this analysis
[3] Confidence interval
[4] Fisher's exact test p-value

**TABLE S6** Descriptive data about the case stables and clinical cases

**TABLE S6a** Frequencies of different paddock surface quality, events, actions, and visits within one month before the onset of the signs in 43 case stables with pastern dermatitis

| Bottom of the paddocks | n[1] | % | 95% CI[2] |
|---|---|---|---|
| Sandy | 23 | 53.5[3] | 38.9–67.5 |
| Wood chips | 1 | 2.3 | 0.4–12.1 |
| Snowy or icy | 32 | 74.4 | 59.8–85.1 |
| Problems in the bottom or legs often muddy | 23 | 53.5[3] | 38.9–67.5 |
| Events, actions, and visits | | | |
| No specific actions | 11 | 25.6 | 14.9–40.29 |
| New horse arrived | 6 | 14.0 | 6.6–27.3 |
| New other animal arrived | 0 | 0 | 0–8.2 |
| Participation in races | 31 | 72.1 | 57.3–83.3 |
| Speed training | 12 | 27.9 | 16.8–42.7 |
| Participation in riding competitions | 3 | 7.0 | 2.4–18.6 |
| Horse visiting outside the stable | 11 | 25.6 | 4.9–40.29 |
| New stable worker started | 1 | 2.3 | 0.4–12.1 |
| New feed lot received | 2 | 4.7 | 1.3–15.5 |
| New bedding lot received | 3 | 7.0 | 2.4–18.6 |
| New training location used | 0 | 0 | 0–8.2 |

[1] Numbers of stables out of all stables in this analysis
[2] Confidence interval
[3] These were not associated with each other (p=0.223)

**TABLE S6b** Combinations of and separate skin lesions in diseased horses in 43 stables with pastern dermatitis

| | n[1] | % | 95% CI[2] |
|---|---|---|---|
| Combinations of skin lesions | | | |
| Ulcers, vesicles, excreting, scabby, oedematous, and erythematous | 18 | 41.9 | 28.4–56.7 |
| Ulcers, scabby, oedematous, and erythematous | 3 | 7.0 | 2.5–18.6 |
| Ulcers, vesicles, excreting, scabby, and oedematous | 3 | 7.0 | 2.5–18.6 |
| Vesicles, excreting, scabby, oedematous, and erythematous | 2 | 4.7 | 1.3–15.5 |
| All other combinations each | 1 | 2.3 | 0.4–12.1 |
| Separate skin lesions | | | |
| Scabby | 41 | 95.3 | 84.5–98.7 |
| Oedematous | 39 | 90.7 | 78.4–96.3 |
| Ulcerative | 37[3] | 88,1 | 75.0–94.8 |
| Erythematous | 35[4] | 87.5 | 73.9–94.5 |
| Excreting | 36 | 83.7 | 70.0–91.9 |
| Vesicular | 28 | 65.1 | 50.2–77.6 |

[1] Number of stables with sign combination or separate sign
[2] Confidence interval
[3] 42 stables answered this question
[4] 40 stables answered this question

**TABLE S6c** Duration of signs and treatments of diseased horses in 43 stables with pastern dermatitis

| | n[1] | mean days | median days | range of days |
|---|---|---|---|---|
| Shortest duration of signs | 39 | 22.3 | 12.0 | from 4 to over 100 |
| Longest duration of signs | 42 | 39.2 | 30.0 | from 2 to over 100 |
| Interval between the onset of signs of the first two cases | 29 | 9.2 | 3.00 | from 0 to 90 |
| Shortest duration of treatment | 31 | 20.5 | 10.0 | from 0 to 120 |
| Longest duration of treatment | 31 | 30.2 | 21.0 | from 0 to 210 |

[1] Numbers of stables that gave answers

**TABLE S6d** Estimation of the responder about the effect of the treatment in diseased horses in 37 stables with pastern dermatitis

| | n[1] | % | 95% CI[2] |
|---|---|---|---|
| Local treatment healed | 27 | 73.0 | 57.0–84.6 |
| Systemic antibiotics[2] healed | 5 | 13.5 | 5.9–28.0 |
| Systemic antibiotics [2] combined to local treatment healed | 1 | 2.7 | 0.5–13.8 |
| Something else healed | 4 | 10.8 | 4.3–24.7 |

[1] Number of stables (6 stables did not answer this question)

[2] Systemic antibiotics were given either perorally or as parenteral injection or both

Note: antibiotics systemically or locally was given to 26 horses (60.5%; 95% CI 45.6-73.6)
